# Multiple types of navigational information are independently encoded in the population activities of the dentate gyrus neurons

**DOI:** 10.1101/2020.06.09.141572

**Authors:** Tomoyuki Murano, Ryuichi Nakajima, Akito Nakao, Nao Hirata, Satoko Amemori, Akira Murakami, Yukiyasu Kamitani, Jun Yamamoto, Tsuyoshi Miyakawa

**Author notes:** **Correspondence:** Tsuyoshi Miyakawa, Division of Systems Medical Science, Institute for Comprehensive Medical Science, Fujita Health University, 1-98 Dengakugakubo, Kutsukake-cho, Toyoake, Aichi 470-1192, Japan.

## Abstract

The dentate gyrus (DG) plays critical roles in cognitive functions such as learning, memory, and spatial coding, and its dysfunction is implicated in various neuropsychiatric disorders. However, it remains largely unknown how information is represented in this region. Here, we recorded neuronal activity in the DG using Ca^2+^ imaging in freely moving mice and analysed this activity using machine learning. The activity patterns of populations of DG neurons enabled us to successfully decode position, speed, and motion direction in an open field as well as current and future location in a T-maze, and each individual neuron was diversely and independently tuned to these multiple information types. In αCaMKII heterozygous knockout mice, which present deficits in spatial remote and working memory, the decoding accuracy of position in the open field and future location in the T-maze were selectively reduced. These results suggest that multiple types of information are independently distributed in DG neurons.

## Introduction

The dentate gyrus (DG) in the hippocampus has been implicated in cognitive functions such as pattern separation^1,2^, contextual encoding^3,4^, and place recognition^3,4,7–9^, and its dysfunction is suggested to be associated with various neuropsychiatric disorders such as epilepsy^10^, schizophrenia^11,12^, bipolar disorder^11,13^, and Alzheimer’s disease^10^. The neural circuitry of DG is composed of excitatory input from layer II neurons in the entorhinal cortex (EC)^14,15^; local interactions among DG granule cells (GCs), mossy cells, and interneurons^4,7,8^; and excitatory output to CA3 pyramidal cells^4,7^. The EC receives multimodal sensory information from other brain regions and contains neurons with distinct functional properties, including grid^16^, border^17^, speed^18,19^, and head-direction cells^20^. Likewise, the hippocampal CA regions receive information from the DG and contains neurons that represent place^21^, locomotion speed^19^, episodic memory^22^, time^23^, and novel spatial experience^24,25^. Therefore, the DG is thought to be involved in the processing and integration of a variety of information^26^.

In spite of its wide-ranging functions, the DG, is unusual among hippocampal regions in that only a small population of principal DG neurons is thought to be active in a given environment^27,28^. In addition, several studies have reported that most DG neurons exhibit poor tuning to position (e.g., DG neurons often have multiple and irregular firing fields^3,29^) and movement speed (e.g., most DG neurons are not specifically tuned to running or resting states^30^). This paradox has led to a significant interest in how information is represented in the population activity patterns of DG neurons^9^.

Here, to unveil important functional characteristics of DG neurons, we performed Ca^2+^ imaging using a micro-endoscope in freely moving mice, which enable us to record activity from a large population of neurons^31,32^. We recorded neural activity of the dorsal DG, that is thought to be relatively more involved in the exploratory behaviour and contextual memory encoding than in ventral DG^5^. Then, we analysed how individual DG neurons are involved in encoding position, speed, and motion direction in an open field as well as current and future location (left or right) in a T-maze. Even if the majority of individual neurons are poorly tuned to these types of information, neurons may encode information by the population activity patterns^9^. Therefore, by using machine learning methods, we also asked if these types of information are encoded in the population activity patterns of DG neurons. Concurrently, to assess how these neural representations might be altered in a disease model, we carried out the same imaging and analysis in heterozygous alpha-calcium/calmodulin-dependent kinase II knockout (αCaMKII^+/−^) mice. Mutations of αCaMKII have been reported to cause intellectual disability accompanied by epilepsy, abnormal emotional and affective behaviour, and/or autistic features^33–36^, and the gene has been also implicated in autism^37^ and bipolar disorder^38,39^. The αCaMKII^+/−^ mice exhibit an array of behavioural abnormalities, including locomotor hyperactivity, impaired working and remote memory, abnormal social/aggressive behaviour, and exaggerated infradian rhythms^39–43^. By studying how neural representation of information is altered in the DG of these mice, we further aimed to gain insight into how multiple types of information are represented in the population activities of dDG neurons in normal and abnormal conditions.

## Results

### 1. Activity patterns of dorsal DG neurons are tuned to the position and speed of the mouse

By using a head-mounted miniaturized microscope, we recorded neural activity in the upper blade of the dorsal DG (dDG) of freely moving mice while they were travelling in an open field (Fig. 1a). We observed that over 90% of all GCaMP expressing neurons in the field of view were active in both of wild-type and αCaMKII^+/−^ mice (Supplementary Fig. 1), and included 72.6±9.9 dDG neurons from 7 wild-type mice and 57.8±9.6 dDG neurons from 5 αCaMKII^+/−^ mice into analysis (for details see “Detection of Ca^2+^ transients” in Materials and Methods section).

**Figure 1.**
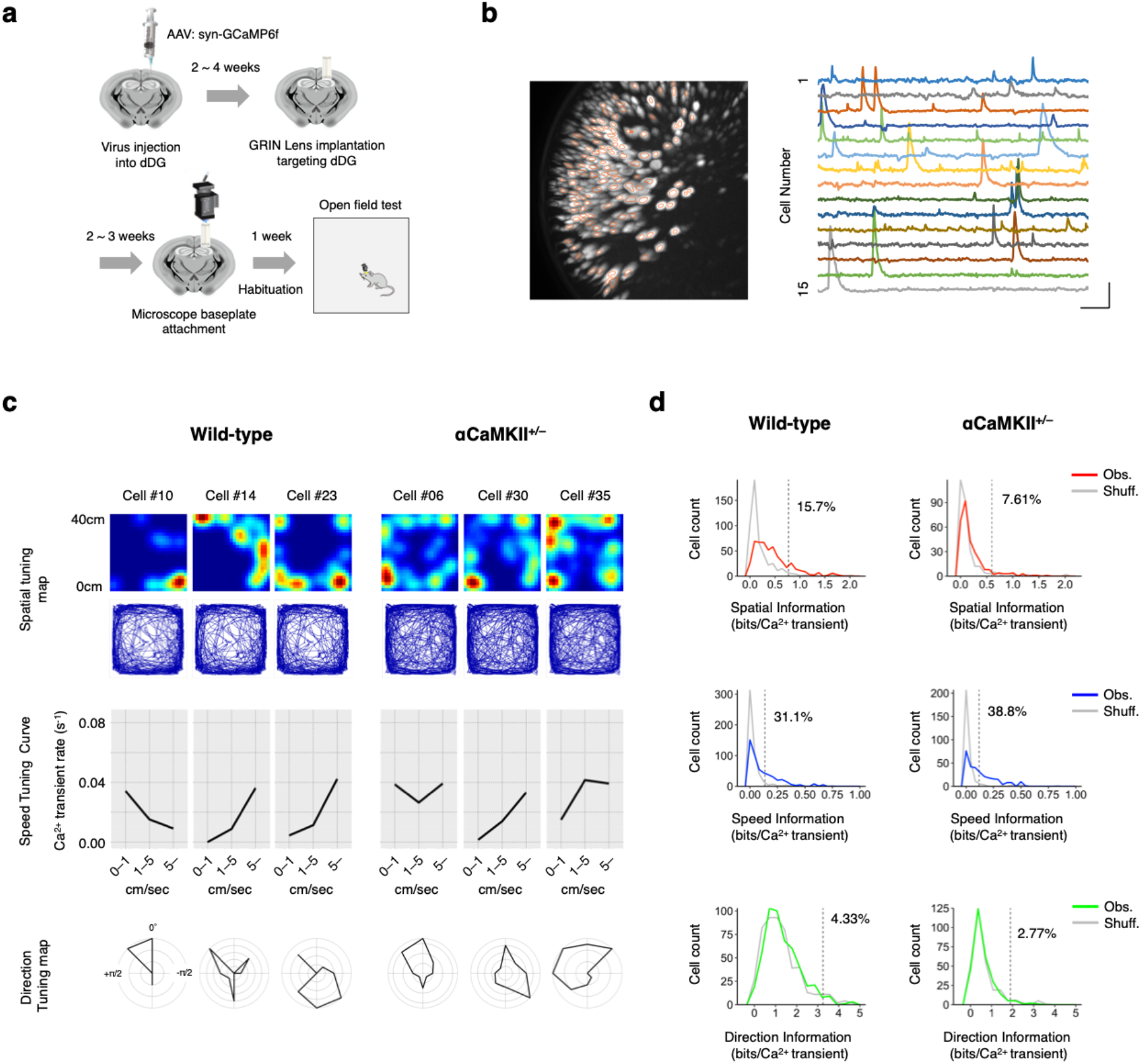
Activity patterns of dorsal DG neurons are slightly tuned to the position and speed of the mouse. **a,** Schematic of the experimental procedures. GCaMP6f was expressed in the dorsal DG (dDG) of mice by AAV injection. At least 2 weeks after viral injection, a graded-index (GRIN) lens was implanted. Two to three weeks after lens implantation, a baseplate for a miniature microscope was attached at the optimal focal plane. Over 1 week after baseplate attachment, the mice were habituated to the test environment and subjected to the behavioural experiments. **b,** Left: Representative image of active dDG neurons in a wild-type mouse. The image shows the projection of the maximum relative fluorescence change (ΔF/F) of the Ca^2+^ transient among the entire set of frames from the recording of the 30 min open field experiment. Orange circles indicate the regions of interest of the recorded dDG neurons. Right: ΔF/F of the Ca^2+^ signals for 15 cells. Scale bars: 30 sec (horizontal) and 5% ΔF/F (vertical). **c,** Colour-coded spatial tuning maps, speed tuning curves, and direction tuning maps of three representative dDG neurons from wild-type mice (left) and αCaMKII^+/−^ mice (right) are shown. Top: Spatial tuning maps showing the density of mouse locations where Ca^2+^ transients were detected. Each map is normalized to each neuron’s maximum Ca^2+^ transient rate and smoothed with a Gaussian kernel (σ = 2.0 cm). Middle: Speed tuning curves representing the mean Ca^2+^ transient rate (s^−1^) when the mouse moved at 0–1 cm/sec, 1–5 cm/sec, and >5 cm/sec. Bottom: Direction tuning maps showing the mean Ca^2+^ transient rate when the motion direction of the mouse was in each of the eight indicated directions. Each map is normalized to the neuron’s maximum mean Ca^2+^ transient rate. North is defined as 0 radians; west is defined from 0 to π radians, and east is defined from 0 to −π radians. **d,** Distributions of information content from observed neurons (“Obs.”, colour-coded curves) and those from shuffled data (“Shuff”, grey curves; 1,000 shuffles per cell; counts normalized by number of shuffles) for all neurons from all animals (508 neurons from 7 wild-type and 289 neurons from 5 αCaMKII^+/−^ mice). The dashed line shows the 95^th^ percentile of the distribution of the shuffled data, and the numbers on the lines represent the percentage of observed neurons exceeding this value.

We analysed the tuning specificities of single dDG neurons regarding mice position, speed, and motion direction. Representative spatial tuning maps, speed tuning curves, and direction tuning maps of individual dDG neurons are shown in Figure 1c. We computed information content (bits/calcium transient)^44^ about the position, speed, and motion direction for each neuron, which we designated spatial, speed, and direction information, respectively. In wild-type mice, the average of spatial information of the dDG neurons was significantly larger than that expected by chance (permutation test, *P*< 10 ^3^) and the effect size was medium (Cohen’s *d* = 0.53). The distribution of the spatial information was shifted to be slightly larger than that of the shuffled data (Fig. 1d), and the significance of spatial information for each cell varied widely from small to large (Supplementary Fig. 2), indicating that most of the dDG neurons are tuned to information about position to varying degrees. The same was also true of the amount of speed information (*P* < 10^−3^; *d* = 0.54; Fig. 1d). The amount of direction information was also larger than that expected by chance (*P* = 3.00 × 10^−3^; Fig. 1d), the effect size was small (*d* = 0.11; Fig. 1d), suggesting that the activity patterns of dDG neurons are relatively weakly tuned to motion direction. Similarly, in αCaMKII^+/−^ mice, the spatial and speed information of dDG neurons were significantly greater than would be expected by chance (spatial, *P* < 10^−3^, *d* = 0.38; speed, *P* < 10^−3^, *d* = 0.58; Fig. 1d), but direction information was not (*P* = 0.03, *d* = 0.01; Fig. 1d). Thus, in both wild-type and αCaMKII^+/−^ mice, individual dDG neurons are tuned to position and speed to varying degrees, but are relatively weakly tuned to motion direction. Indeed, it appears that these information types are widely distributed in the populations of dDG neurons.

### 2. Information about position, speed, and motion direction is encoded in the population activity patterns of dDG neurons

Next, to examine the information coding at the population level, we tested whether position, speed, and motion direction of the mice could be decoded from the population activity of dDG neurons using machine learning methods. Eight models were considered for decoding analysis in this study, and since there was no significant difference in the decoding accuracy among these models, we used the LSTM (Long Short-Term Memory), which showed a slightly higher accuracy than others (Supplementary Fig. 3). Representative decoding results are shown together with the actual positions, speeds, and motion directions in Figure 2a–c (Decoding errors for all mice are shown in Supplementary Fig. 4-6). In wild-type mice, the decoding accuracies for position, speed, and motion direction were significantly higher than those expected by chance, which is the decoding accuracy when using the shuffled data (paired *t*-test, position, *P* = 2.18 × 10^−4^; speed, *P* = 2.42 × 10^−3^; motion direction, *P* = 1.36 × 10^−3^; Fig. 2d-f, left panels). These results indicate that information about position, speed, and motion direction are encoded in the population activity of dDG neurons in wild-type mice.

**Figure 2.**
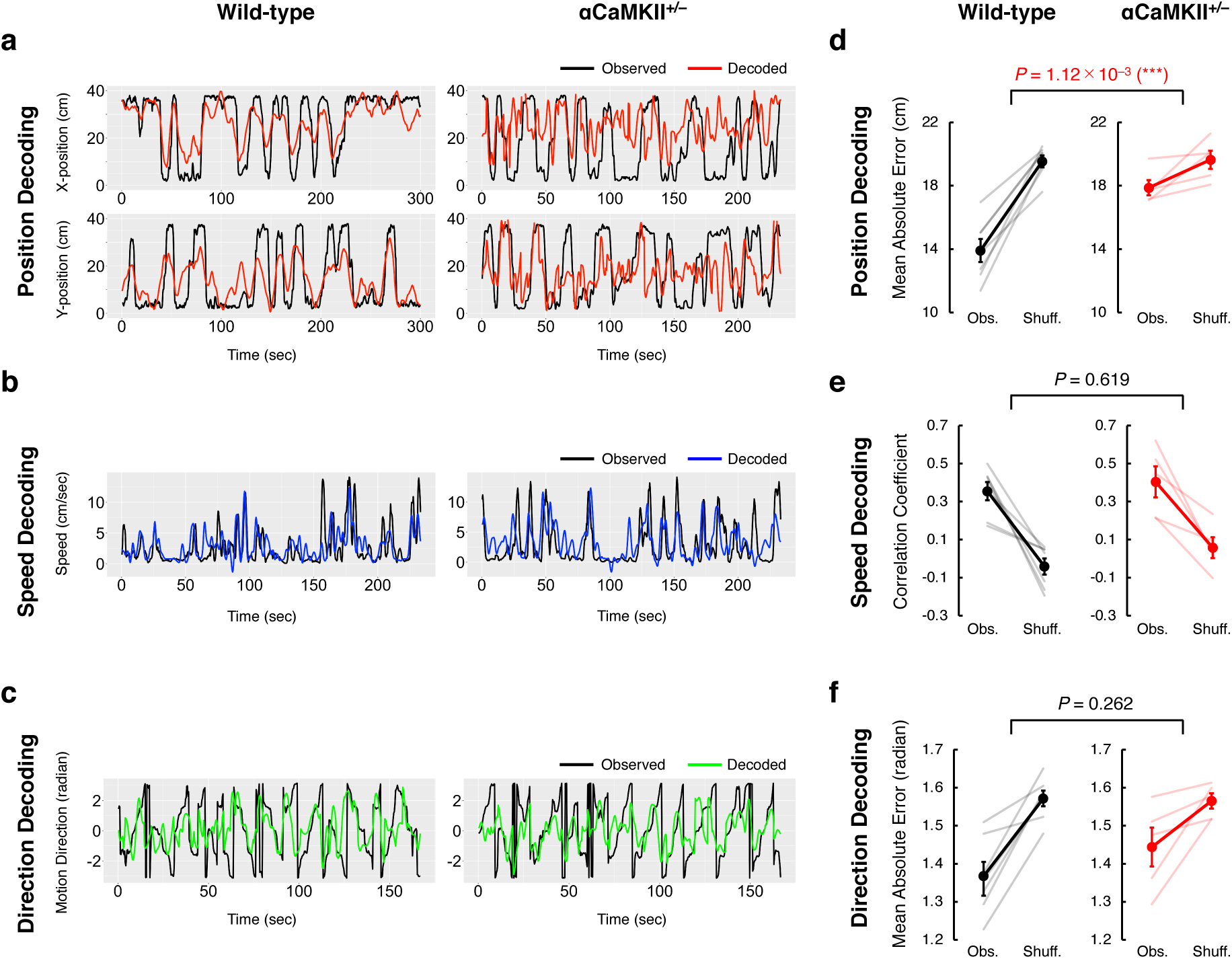
Information about position, speed, and motion direction is encoded in the population activity patterns of dDG neurons. **a,** Representative results of position decoding from one mouse each for wild-type mice (left panels) and αCaMKII^+/−^ mice (right panels). The X and Y positions (cm) of the mouse in the open field are shown in the upper and lower panels, respectively. Black lines show the observed X and Y positions of the mice (observed), and the red lines show the positions decoded from the population activity patterns of dDG neurons (decoded). **b,** As for panel a, but for speed instead of position. **c,** As for panel a, but for direction instead of position. **d,** Accuracy of position decoding in 7 wild-type mice (left) and 5 αCaMKII^+/−^ mice (right). The accuracy of position decoding is reported as the mean absolute error. Black and red lines represent means, and error bars indicate the standard error of the mean (SEM). **e,** Accuracy of speed decoding, represented as the correlation coefficient between the observed and decoded speeds of the mice in each time bin. **f,**Accuracy of direction decoding. The decoding error for motion direction is computed as mean absolute error between the observed and decoded motion directions in each time bin.

In αCaMKII^+/-^ mice, the decoding accuracies for speed and motion direction were significantly higher than those expected by chance (paired *t*-test, speed, *P* = 1.66 × 10^−2^; motion direction, *P* = 0.0210), and not significantly different from those of wild-type mice (unpaired *t*-test, Obs.-Shuff. is compared; speed, *P* = 0.619; motion direction, *P* = 0.262; Fig. 2e, 2f). On the other hand, the decoding accuracy of position was better than chance to a marginally significant degree (*P* = 0.0549), but was significantly lower than that of wild-type mice (unpaired *t*-test, Obs.-Shuff. is compared; *P* = 1.12 × 10^−3^; Fig. 2d). This lower position decoding accuracy in αCaMKII^+/-^ mice compared to wild-type mice might not be due to the difference in the number of the neurons used for decoding between them (position decoding accuracy × the number of neurons, *R* = −0.0279; Supplementary Fig. 7). Thus, in αCaVIKII^+/-^ mice, position information is selectively impaired in the dDG, whereas speed and motion direction information is not. These results imply that with respect to information about speed and motion direction, information about position is encoded in the dDG by a different population of neurons, a different coding principle, or both.

### 3. Information about position, speed, and motion direction is independently distributed in the population activity patterns of dDG neurons

Here, we investigated how each neuron encodes position, speed, and motion direction. One possibility is that a neuron may carry a particular type of information exclusively. Another possibility is that a neuron may carry multiple types of information. In this case, some particular groups of neurons may carry a relatively larger amount of information about two or more types of information than others, or the amount of one type of information carried by a neuron may be independent from that of other types of information.

At first, to clarify the relationship among the amount of information encoded by individual neurons for these three information types, we examined correlations between them. Scatter plots for each pair of variables (among position, speed, and direction information) show fairly uniform distributions and relatively weak correlations (spatial × speed, *R* = 0.167; speed × direction, *R* = −1.67 × 10^−4^; direction × spatial, *R* = 0.257; Fig. 3a). These weak correlations suggest that there is limited correspondence between how each form of information is encoded in individual neurons.

**Figure 3.**
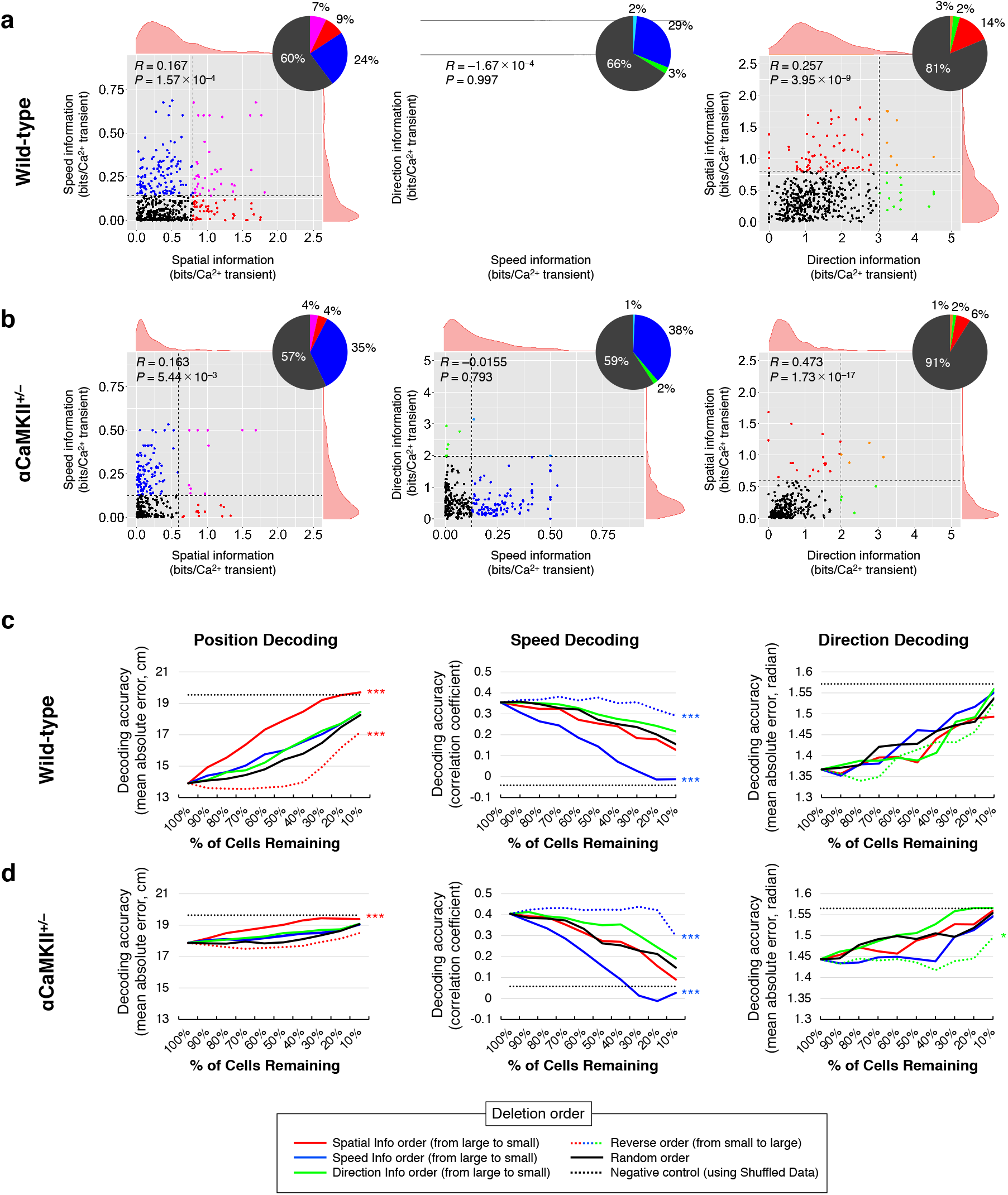
Information about position, speed, and motion direction is independently distributed in the population activity patterns of dDG neurons. **a,** Left panel: Scatter plot showing the distribution of spatial and speed information from 508 dDG neurons from 7 wild-type mice. Each dot corresponds to an individual neuron. Vertical and horizontal dashed lines indicate the 95^th^ percentile of spatial and speed information obtained from shuffled data (same as in Fig. 1d). Pie chart represents the percentage of neurons with spatial and speed information above or below this value. *R* values are correlation coefficients. Middle and right panels: Scatter plots showing the distribution of speed and direction information and of direction and spatial information, respectively. **b,** As for panel a, but for 289 dDG neurons from 5 αCaMKII^+/−^ mice. **c,** Changes in decoding performance when neurons are individually deleted from the datasets (left; position decoding, middle; speed decoding, right; direction decoding). Each line shows the average data from the 7 wild-type mice (\**P* < 0.05, \*\**P* < 0.01, \*\*\**P* < 0.001; Two-way ANOVA with Tukey-Kramer post hoc tests, vs random order deletion). **d,**As for panel c, but showing average data from the 5 αCaMKII^+/−^ mice.

Next, we examined whether populations of dDG neurons used for coding position, speed, and motion direction are exclusive from, identical to, or independent of each other, by assessing how the decoding performance changes when neurons are deleted from the datasets one by one. In our analysis, when neurons were deleted from the dataset in order of decreasing spatial information, position decoding accuracy decreased more rapidly than when they were deleted randomly (Two-way ANOVA with Tukey-Kramer post hoc tests, Spatial info order × Random order, *P* = 2.29 × 10^−10^; Fig. 3c, left panel); the reverse was true of deletion in order of increasing spatial information (Reverse spatial info order × Random order, *P* = 5.20 × 10^−4^; Fig. 3c, left panel). These results indicate that neuron populations consisting of individual neurons with larger amounts of spatial information tend to be more important in the population coding of position information than those with smaller amount of spatial information. Furthermore, the removal of neurons in decreasing speed or direction information order mimicked the impacts of randomly ordered deletion on position decoding performance (Fig. 3c, left panel). These results support the notion that the information about position is independently encoded from that of speed or motion direction. Similarly, the removal of neurons in decreasing speed information order lowered speed decoding accuracy to a greater extent than randomly ordered deletion (Speed info order × Random order, *P* = 1.40 × 10^−8^; Fig. 3c, middle panel); on the other hand, deletion in the reverse order had smaller impacts on speed decoding accuracy (Reverse speed info order × Random order, *P* = 4.23 × 10^−3^; Fig. 3c, middle panel). Removal of neurons in increasing spatial or direction information order did not decrease speed decoding performance more rapidly than randomly ordered deletion (Fig. 3c, middle panel), showing that the amount of speed information in a neuron is also independent from the amount of spatial or direction information. For direction decoding, there was no significant difference in the decoding accuracy for the removal of neurons in decreasing spatial/speed/direction information order, increasing direction information order, or random order (Fig. 3c, right panel). These results showed that the amount of spatial, speed, or direction information in an individual neuron is not associated with its contribution to the decoding accuracy of motion direction. Thus, neuron populations involved in coding these three types of information are not mutually exclusive or identical to each other, but rather are independent of each other.

Furthermore, as described above, there is another possibility that different types of information are encoded in the dDG by a different coding principle (e.g., in CA1 and EC, position information is represented in the activity patterns of neurons^3,45^, and speed information is encoded by the frequency of the neural activity^18,19^). Then, at first, we examined the relationship between position decoding accuracy and distinctness of activity patterns of dDG neurons across the subareas in the open field. In our data, the population vector overlap (PVO) (see Materials and Methods section; Supplementary Fig. 8a), which measures how similar firing patterns are across different subareas in the open field, was significantly correlated with the error of position decoding (Supplementary Fig. 8b). Thus, the increased position decoding error in αCaMKII^+/-^ mice relative to that in wild-type mice (*P* = 1.12 × 10^−3^, Fig. 2d) might be attributable to the larger mean PVO (*P* = 0.0208, Supplementary Fig. 8b), corresponding to less distinct firing patterns of neural ensembles across different subareas. Next, we investigated how encoding of speed information is associated with overall firing frequency of dDG neurons. We found positive and linear correlations between the speed of the mice and the average Ca^2+^ transient rate in the dDG (Supplementary Fig. 8c). We also found a significant correlation between speed decoding accuracy and the change in the average Ca^2+^ transient rate with speed (Supplementary Fig. 8d), neither of which were significantly different between wild-type and αCaMKII^+/-^ mice (Fig. 2e, Supplementary Fig. 8d). Thus, information about position and speed might be encoded with different coding principles in the dDG, and only the coding principle for position may be affected in the dDG of αCaMKII^+/-^ mice.

### 4. The activity patterns of dDG neurons are tuned to current and future locations to varying degrees

Next, we recorded the activity patterns of dDG neurons while the mice performed a forced-alternation task in a modified T-maze (Fig. 4a; for details, see “Behavioural experiments” in Materials and Methods section). We used the T-maze test because it has been used previously to assess working memory in rodents^46^ and because dysfunction in the DG is associated with reduced performance in this test^46^. Wild-type mice made the correct choice (that is, during the free-choice period, they chose the arm opposite the arm that was open in the forced-choice period) 71.4% of the time, averaged of 50 sessions. In contrast, the αCaMKII^+/-^ mice showed 52.0% correct choice on the average, which is close to chance (50%) and significantly lower than the performance of the wild-type mice (*P* = 2.38 × 10^−3^; Fig. 4b), indicating that the αCaMKII^+/−^ mice showed deficits in this working memory task.

**Figure 4.**
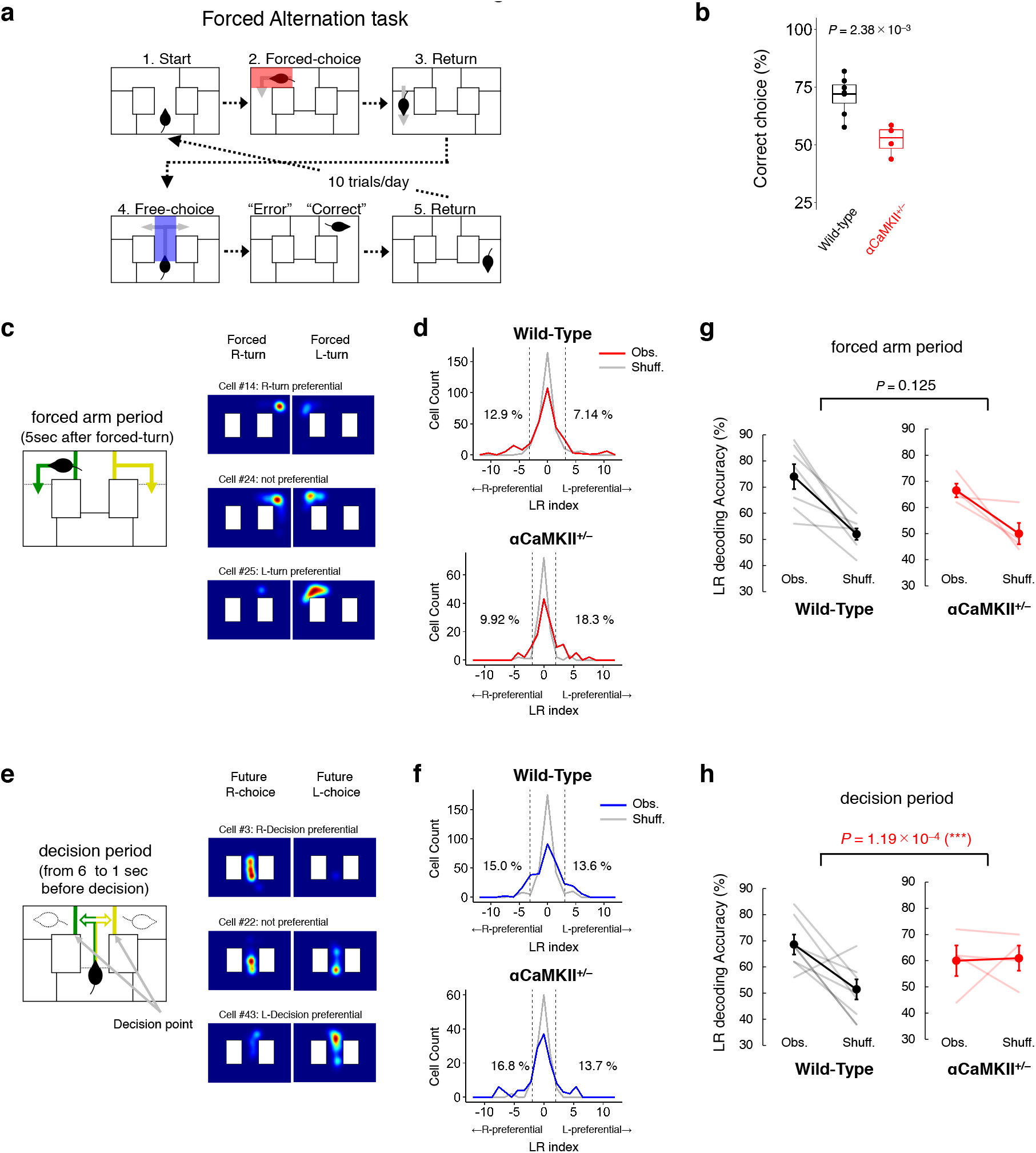
Information about current and future LR location is encoded in the population activity patterns of dDG neurons. **a,** Forced-alternation task in a modified T-maze test. Each trial consists of a forced-choice run followed by a free-choice run. In the forced-choice run, the mouse is forced to turn one of two randomly selected directions (1. Start→2. Forced choice), whereas in the free-choice run, the mouse can choose to turn left or right (3. Return→4. Free choice). This task exploits the behavioural tendency of mice to choose the arm opposite the one previously chosen. Therefore, if the direction that the mouse chooses in the free-choice run is opposite to the direction presented to it in the forced-choice run, the trial is considered “correct”; otherwise, it is considered an “error”. **b,** The correct choice rate (%) in the forced-alternation task in 7 wild-type mice and 4 αCaMKII^+/-^ mice. The centre lines, box boundaries and whiskers for the boxplot indicate median, upper and lower quartile, and maximum and minimum of the data. **c,** The forced arm period was defined as the 5 sec period after the mouse performed the forced turn (left). Color-coded cell activity maps of three representative dDG neurons from wild-type mice (right). These maps show the density of mouse locations where Ca^2+^ transients were detected in the cases of L-turn or R-turn in all 50 trials. Each map is normalized to each neuron’s maximum Ca^2+^ transient rate and smoothed with a Gaussian kernel (σ = 2.0 cm). **d,** Distributions of LR indices calculated from the recordings of the observed neurons (red curve, “Obs”) and from shuffled data (grey curve, “Shuff.”; 1000 shuffles per cell; counts normalized by number of shuffles) in the arm period, which corresponds to the 5 sec following the forced turn. All 294 neurons from 7 wild-type mice and 131 neurons from 4 αCaMKII^+/-^ mice are shown in histograms. The dashed line shows the 95^th^ percentile of the distribution of the shuffled data, and the numbers on the lines represent the percentage of observed neurons exceeding this value. **e,** The decision period was defined as the 6 to 1 sec period prior to the free-choice turn (left). As for panel (c), but for color-coded cell activity maps of representative dDG neurons in the decision period (right). **f,** As for panel (d), but for LR indices in decision period, which corresponds to the 6 to 1 sec before the turn decision is made. **g,** LR decoding accuracy in wild-type mice and αCaMKII^+/-^ mice from the population activity patterns of dDG neurons in the arm period. The grey (left panel, wild-type mice) and pale red lines (right panel, αCaMKII^+/-^ mice) show the decoding accuracy of the individual mice in each group. Black (left panel, wild-type mice) and red (right panel, αCaMKII^+/-^ mice) lines indicate the average decoding accuracy of all mice in each group. Error bars indicate SEM. **h,**As for panel (e), but for the decision period.

Next, to elucidate the functional properties of dDG neurons in T-maze test, we calculated the “LR index”, which takes a higher absolute value if a neuron shows differential activity patterns when the mouse is on either the left or right side of the apparatus. Here, “current location” designates the (left or right) location of the mouse at time of recording. These neural activity patterns were taken from 5 sec after the mouse performed the initial forced turn, which was designated the “forced arm period” (Fig. 4c). “Future location” refers to the (left or right) location of the mouse after the “decision period”, during which the neural activity patterns for calculating the LR index and decoding the future LR location were taken. The “decision period” was defined as the period from 6 to 1 sec prior to the free-choice turn (Fig. 4e). Note that in this period, the future location did not significantly correlate with physical location, speed, and motion direction of the mouse (Supplementary Fig. 9g–i). In the wild-type mice, the absolute values of the LR indices of neurons during the forced arm period were significantly higher than those expected by chance (permutation test, *P* < 10^−3^) and the effect size was medium (Cohen’s *d* = 0.43). The distribution of LR indices of neurons was shifted to be slightly larger in absolute values than that of the shuffled data (Fig. 4d, upper panel), showing that the activity patterns of dDG neurons are tuned to left or right of the current location to varying degrees. The same was also true of the LR indices of neurons during the decision period in wild-type mice (*P* < 10^−3^, *d* = 0.52; Fig. 4f, upper panel) and of those in the forced arm and decision period in αCaMKII^+/−^ mice (forced arm period, *P* < 10^−3^, *d* = 0.51; decision period, *P* < 10^−3^, *d* = 0.54; Fig. 4d and 4f, lower panel). These results showed that the activity patterns of dDG neurons of wild-type and αCaMKII^+/−^ mice are tuned to both current and future LR locations to varying degrees.

### 5. Information about current and future LR location is encoded in the population activity patterns of dDG neurons

Next, we trained a binary decoder on the population activity data obtained from dDG neurons and tested whether it was able to estimate the LR location of the mice in the forced arm and decision periods. In wild-type mice, we were able to decode the current LR location of the mice in the forced arm period from the population activity patterns of dDG neurons more accurately than those expected by chance (paired *t*-test, *P* = 1.08 × 10^−3^; Fig. 4g, left panel). We were also able to estimate the future LR location of mice from population activity patterns in the decision period more accurately than that expected by chance (*P* = 0.0117; Fig. 4h, left panels). These results suggest that information about the current and future LR location may be encoded in dDG neuron activity patterns obtained in the current and decision period, respectively. Meanwhile, in αCaMKII^+/-^ mice, the decoding accuracy of the current LR location in the forced arm period was more accurate than that expected by chance (*P* = 0.0220; Fig. 4g, right panel) and not significantly different from that of wild-type mice (unpaired *t*-test, Obs-Shuf. is compared, *P* = 0.125; Fig. 4g), indicating that information about current LR location is encoded in dDG neuron activity patterns of αCaMKII^+/-^ mice. We also aimed to estimate the future LR location of αCaMKII^+/-^ mice from the population activity of the decision period, but its accuracy was not higher than that expected by chance (*P* = 0.873; Fig. 4h, right panel) and was significantly lower than that of wild-type mice (*P* = 1.19 × 10^−4^; Fig. 4h), suggesting that future LR choice is not accurately represented in dDG neurons in αCaMKII^+/-^ mice. Thus, the neural representation of future location is selectively impaired in the dDG of αCaMKII^+/-^ mice, whereas that of the current location is not, implying that current and future locations in the T-maze may be independently encoded among the populations of dDG neurons.

### 6. Information about current and future LR locations is widely distributed in the population activity patterns of dDG neurons

Next, we investigated the relationship between LR indices in forced arm and decision periods of each dDG neurons. The scatter plot of the LR indices of the dDG neurons obtained in the forced arm and decision periods of the wild-type mice showed a small but statistically significant correlation (*R* = 0.2281, *P* = 7.91 × 10^−5^; Fig. 5a). Meanwhile, in αCaMKII^+/-^ mice, the LR indices of the neurons obtained in the forced arm and decision periods showed a uniform distribution, and these indices were not significantly correlated (*R* = −0.1210, *P* = 0.1687; Fig. 5b). Thus, the LR-tunings of dDG neurons about current and future LR locations are weakly correlated in wild-type mice and largely independent of each other in αCaMKII^+/-^ mice. These correlations suggest that there is a significant, but weak, relationship between LR-tunings of individual dDG neurons during the forced arm- and decision-period in wild-type mice.

**Figure 5.**
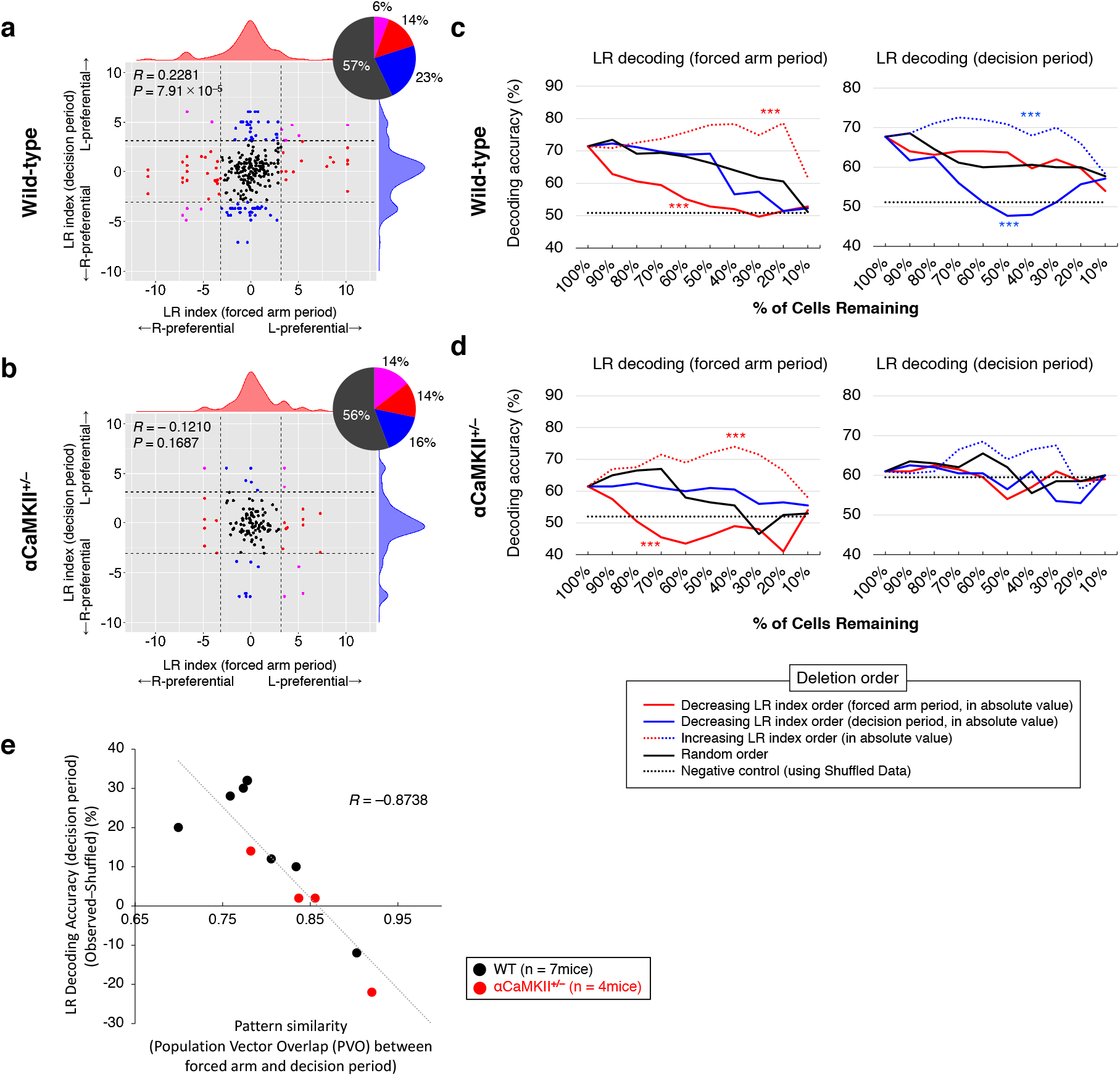
Information about current and future LR locations is widely distributed in the population activity patterns of dDG neurons. **a, b,** Scatter plot showing the distribution of the LR indices of dDG neurons in the forced arm and decision periods. Each dot corresponds to an individual neuron. A total of 294 neurons from 7 wild-type mice (a) and 131 neurons from 4 αCaMKII^+/-^ mice (b) are shown. The horizontal and vertical axes indicate the LR indices from the forced arm and decision periods, respectively. Vertical and horizontal dashed lines indicate the 95^th^ percentile of LR indices from the forced arm and decision periods obtained from shuffled data (same as in Fig. 4c, 4e). Pie charts represent the percentage of neurons with LR indices above or below this value. *R* values are correlation coefficients. **c, d,** Changes in LR decoding performance (left panels, arm period; right panels, decision period) when neurons were individually deleted from the datasets for wild-type mice (c) and αCaMKII^+/-^ mice (d). The horizontal axis represents the percentage of remaining neurons that are used for population decoding after neurons are deleted. The vertical axis shows the LR decoding accuracy (%) in the arm period (left panels) and the decision period (right panels). Each line shows the average of 7 wild-type mice (c) and 4 αCaMKII^+/-^ mice (d) (\**P* < 0.05, *\*\*P* < 0.01, *\*\*\*P* < 0.001; Two-way ANOVA with Tukey-Kramer post hoc tests, vs random order deletion). **e,** Scatter plot showing the correlation between the PVO (forced arm period × decision period) and LR decoding accuracy in the decision period, which is computed as the decoding accuracy obtained from observed data minus those of shuttled data (Obs–Shuffled, %). The individual black and red dots correspond to individual wild-type and αCaMKII^+/-^ mice, respectively.

To examine how information about the current and future LR locations is distributed in the population activity patterns of dDG neurons, we assessed how the LR decoding accuracy in the forced arm and decision periods changed when neurons were removed from the datasets one by one. In the data collected in wild-type mice during the arm period, upon deletion of neurons in order of decreasing LR index for the forced arm period, the LR decoding accuracy of the forced arm period decreased more rapidly than it did upon randomly ordered deletion (Two-way ANOVA with Tukey-Kramer post hoc tests, *P* = 1.54 × 10^−6^; Fig. 5c, left panel); deletion of neurons in order of increasing LR index had smaller impacts on the LR decoding accuracy than randomly ordered deletion (*P* = 1.12 × 10^−5^; Fig. 5c, left panel). However, the removal of neurons in order of increasing LR index for the decision period did not decrease decoding accuracy more rapidly than randomly ordered deletion (Fig. 5c, left panel). Meanwhile, for LR decoding in the decision period, the deletion of neurons in order of increasing LR index for the decision period order significantly decreased the decoding accuracies (*P* = 1.92 × 10^−2^; Fig. 5c, right panel), and reverse order deletion showed smaller decreases than that by random order deletion (*P* = 1.27 × 10^−2^; Fig. 5c, right panel). On the other hand, the removal of neurons in order of decreasing LR index for the forced arm period did not have much impact on LR decoding performance, similar to the impact of randomly ordered deletion (Fig. 5c, right panel). These results suggest that neuron population involved in coding current location is independent from that of future location, and vice versa. Thus, information about the current and future location is expected to be independently distributed in the population activity patterns of dDG neurons.

As described above, we noted a small but significant correlation between the LR indices of the forced arm and decision periods in wild-type mice (*R* = 0.2281, *P* = 7.91 × 10^−5^; Fig. 5a). We sought to examine whether associations between neural activity patterns during the forced arm and decision periods correlate with an accurate neural representation of future LR location during the decision period. First, we calculated the PVO (see Materials and Methods section) of the averaged neuronal activity frequency between the forced arm and decision periods for each mouse. We next compared them with LR decoding accuracy during the decision period. We here defined decoding accuracy as the difference between observed and shuffled data (Obs–Shuff) to standardize, because decoding accuracy expected by chance in decision period can be larger than 50% due to the bias of LR choice of mouse (Fig. 4h). Our result showed that there was a significant negative correlation (*R* = −0.8738; Fig. 5e), showing that the LR decoding accuracy during the decision period was higher, PVO tend to be smaller, meaning that neuronal activity patterns in these periods are in more contrasting patterns (i.e., if a neuron is relatively active in one period, it becomes relatively inactive in another period). These results indicate that information about future LR location is more accurately represented in the dDG when there is a stronger association and contrasting patterns between the population activity of dDG neurons in the forced arm and decision periods.

## Discussion

In this study, we demonstrated that multiple types of information (position, speed, and motion direction in an open field as well as current and future left or right location in a T-maze) were represented in the population activity patterns of dDG neurons. These multiple types of information were encoded by overlapping but independent populations of dDG neurons. Furthermore, in αCaMKII^+/-^ mice, which present deficits in spatial remote and working memory, the neural representation of information about position in the open field and future LR location in the T-maze was selectively impaired, supporting the notion that different types of information are independently distributed among dDG neurons.

The accurate decoding of position (in the open field) and current LR location (in the T-maze) in wild-type mice indicates that the DG is involved in place coding, which is consistent with previous studies^3,4,7,8^. Additionally, the successful decoding of speed and motion direction in the open field and future location in the T-maze in this study indicates that the dDG is also involved in the neural representation of these types of information, which only a few studies have reported^30,9^. It is of interest that the decoding accuracies in our study were accurate despite the DG neurons is generally thought to have a lower range of firing frequencies compared to that of most other brain regions^28^. Recently, recordings using tetrode^7^, silicon probe^4^, and calcium imaging^8^ showed that the activity patterns of GCs were at much lower frequency than those of DG mossy cells and CA3 pyramidal cells^4,7,8,28^. In our data, the average recorded Ca^2+^ transient frequency was quite low, approximately 0.02 Hz, which was comparable to that of GCs (1–2/min) reported by the previous imaging study^8^. Our results thus suggest that the dDG neurons may be involved in information coding with their low-frequency activity patterns.

We found that the populations of DG neurons cooperate to encode information about position, speed, and motion direction in an open field, indicating that these types of information are diffusely distributed among dDG neurons, which is consistent with a recent report^9^. In this report, the authors proposed that the interpretability of individual neurons is not necessarily important for their contribution to information coding based on the weak correlation between an index of spatial information and the “importance index”, a measure of the contribution of each neuron to decoding. We evaluated the importance of a neuron in decoding by assessing how the decoding performance changes when a neuron is deleted from the population. In contrast with their conclusion, our results indicate that dDG neurons with a larger amount of information are more important for decoding position and speed in an open field than those with smaller amounts of information. In addition, we identified redundancy in the distribution of some types of information in the dDG; our results indicate that only the top 30-40% of all neurons were sufficient for decoding position and speed with maximum accuracy (Fig. 3c; red dotted line in the left panel; blue dotted line in the middle panel). On the other hand, approximately 70-80% of all the recorded neurons can contribute to the accuracy of spatial or speed decoding (Fig. 3c; solid red line in the left panel; solid blue line in the middle panel). The neurons with an intermediate information content, which are not needed for population encoding when the neurons with the largest information content are spared, can contribute to the decoding accuracy when these top-class neurons were deleted, further demonstrating the redundancy of information encoding in the individually recorded neurons. Furthermore, although we recorded only approximately 70-80 neurons on average per wild-type mouse, out of approximately 500,000 neurons in the entire DG^47^, our decoding, based on the activity patterns of this small number of cells, was fairly accurate. It has been reported that there is a difference in the distribution of functional cell types between the dorsal and ventral DG^5^; even taking this into account, it is assumed that there are far more neurons in the entire dorsal DG than are needed to accurately encode these types of information. Generally, the number of states that a neural circuit can represent is thought to depend on the number of neurons and the dynamic range of firing frequencies of the neurons^28^. Given this assumption, we propose that the diffuse, redundant encoding of information in a large number of infrequently firing neurons allows the DG to represent a large number of behavioural states in its population activity patterns.

Remarkably, we also discovered that some of the neurons of the DG demonstrate multiplexity, i.e., each of these neurons is involved in encoding multiple types of information. Neuronal multiplexity has also been reported in other brain regions, including the medial EC, in which some neurons can encode position, speed, and head direction^48^, and the parietal and frontal cortexes, in which some neurons can encode multiple behavioural or task-relevant variables^49,50^. Neuronal multiplexities are thought to be essential for representing a large number of independent variables in the population activity patterns of a limited number of neurons^49,50^. It is known that if the functional tunings for multiple variables in an individual neuron are in patterns similar to those of other neurons, the number of independent variables that can be represented by the population activity patterns will be reduced. This is because the dimensions represented by the collective activity of these neurons would not be larger than that of a single neuron^50^. In other words, to represent many types of information, it is necessary for neurons to be tuned to different types of information in patterns that are independent of each other. While a previous report proposed that the groups of neurons encoding position partially overlap with those encoding motion direction^9^, the dependency/independency of the distribution of these types of information was not evaluated. In our study, we found that multiple types of information are independently distributed among dDG neurons by quantitatively evaluating the importance of the neurons involved in the decoding (Fig. 3c). The selective impairments of information (position in open field and future location in T-maze) in αCaMKII^+/−^ mice also support our claim that multiple types of information are independently distributed. The multiplexity and independence of dDG neurons that we have demonstrated in this study suggests that the involvement of dDG neurons in encoding certain types of information does not prevent them from encoding other types of information. Together with their diffuseness and redundancy, the multiplexity and independence of dDG neurons seem to allow the DG not only to express a large number of states for each type of information but also to express these states for multiple types of information. Other than the information types described in this study, some other types of information are thought to be encoded in the DG, such as vestibular, olfactory, visual, and auditory information^26,51^, which may also be encoded by taking advantage of the diffuseness, redundancy, and multiplexity of DG neurons.

Mutations of αCaMKII have been reported to cause intellectual disability accompanied by epilepsy, abnormal emotional and affective behaviour, and/or autistic features^33–36^, and the gene has also been implicated in autism^37^ and bipolar disorder^38,39^. αCaMKII^+/−^ mice have an endophenotype called “immature dentate gyrus (iDG)”, in which the neurons in the dentate gyrus are in a pseudo-immature status^39,42^. The phenomenon similar to their iDG can be seen in other mouse models of neuropsychiatric disorders and human patients^43,52^. A prominent feature of this phenotype is that the induction of expression of immediately early genes, such as c-Fos and Arc, are almost completely absent in the granule cells in the DG^42,53–55^. Interestingly, despite a dramatic reduction in the expression of c-Fos and Arc, the DG neurons of αCaMKII^+/−^ mice seem to fire in a fairly normal manner with regard to the frequency of calcium transients (Supplementary Fig. 1), indicating that the machinery linking calcium transients and the expression of those genes is disrupted in this phenotype and that the expression of c-Fos or Arc is not always a good index of neuronal firing. Another interesting result obtained from the αCaMKII^+/−^ mice was that the decoding accuracy of position in the open field was selectively impaired in the dDG, while those of speed and direction were not. This selective impairment is unlikely to be attributed to selective death or general functional impairment of specific groups of neurons because we found that neurons carrying position information also encode other types of information in wild-type mice. Instead, we assume that different types of information are encoded by different coding principles and that the coding principle for position information is selectively impaired in the dDG of αCaMKII^+/−^ mice. Previous studies have reported that successful spatial coding relies on having distinct firing patterns of hippocampal neurons for different locations^3,45^. Our results showed that, similar to the hippocampus, distinct firing patterns across different subareas in the dDG are significantly associated with accurate spatial coding, suggesting that position information might be encoded by the ensemble activities of neurons. On the other hand, it has been reported that neurons in the hippocampus and EC show positive and largely linear firing responses to running speed^18,19^. In our data, we also found a significant correlation between speed decoding accuracy and the average Ca^2+^ transient rate. Thus, information about position and speed might be encoded with different coding principles in the dDG, and only the coding principle for position may be affected in the dDG of αCaMKII^+/−^ mice. Furthermore, it was recently reported that the cell death of interneurons between the CA1 region and the DG in epileptic mice causes desynchronized hippocampal activity, which results in poor spatial processing in the hippocampus^45^. It is also known that hippocampal pyramidal neurons of mice with a point mutation in the αCaMKII show a high firing rate and poor spatial selectivity^56^. Increased excitability and epileptiform activity have also been observed in the DG and CA1 of mice with mutations in αCaMKII^+/−^, which may lead to disorganized activity patterns in the neurons of these regions^57,56,42,58^. Thus, the selective impairment of position information in the DG of αCaMKII^+/−^ mice may be attributed to an increased excitability and disruption of the organized activity patterns of its neurons. On the other hand, speed information in the dDG of αCaMKII^+/−^ mice was not disturbed, possibly because it depends on changes in the neural firing frequency, which is not affected by increased neuronal excitability. The dDG may utilize different coding principles for encoding different types of information, which supports the notion that individual neurons can independently participate in encoding different types of information. Future studies on the mechanism underlying the selective impairment of information in αCaMKII^+/−^ mice and other model mice that show similar phenotypes would contribute to the understanding of the pathophysiology of neuropsychiatric disorders that share this phenotype.

The traditional view of place coding in the hippocampal circuit is that the place cells encode the current location of the animal. It has also been known that neural activity patterns in CA1^25^, CA3^59^, and medial EC^60^ are involved in the prospective representation of future states. In our analysis, we were able to decode whether the mice were on the left or right side of the T-maze by the population activity patterns of dDG neurons either after or before a left or right turn. These results suggest that the population activity patterns of dDG neurons not only encode the current location, but also may be involved in the predictive representation of future states, alongside other hippocampal regions. Furthermore, our result in Fig. 5e showed that this predictive representation of the future states seem to be formed in the population activity pattern of dDG neurons, especially when they are associated with those of the recent past.

In conclusion, our findings suggest that multiple types of information are diffusely, redundantly, and independently encoded in the population activity patterns of dDG neurons. These features of information coding may be the basis on which the DG is involved in the processing and integration of variety of information. Future studies are needed to determine whether information about other functions of the DG (e.g., episodic memory, object recognition, and odour information processing) is similarly encoded in the population activity patterns of neurons, and how this encoding is altered in disease states (e.g., amnesia, Alzheimer disease, and schizophrenia).

## Materials and Methods

### Animals

All experimental protocols were approved by the Institutional Animal Care and Use Committee of Fujita Health University. Adult male C57BL/6J mice and αCaMKII^+/-^ mice were obtained from Jackson Laboratories (Bar Harbor, Maine) and were backcrossed to C57BL/6J mice (Charles River, Yokohama) for at least 19 generations. They were used for experiments at 50 ± 4.4 weeks of age. The mice were housed one per cage in a room with a 12 hr light/dark cycle (lights on at 7:00 a.m., off at 7:00 p.m.) with access to food and water *ad libitum*. The room temperature was maintained at 23 ± 2 °C. All experiments were conducted during the light period.

### Surgeries

#### Viral delivery of Ca^2+^ sensor

For delivery of a fluorescent Ca^2+^ sensor into dorsal DG neurons, adeno-associated virus (AAV) carrying the GCaMP6f vector was injected into the DG of the dorsal hippocampus in adult mice by a conventional method (right hemisphere; 2.0 mm posterior to bregma (AP), 1.0 mm lateral to midline (ML), and 2.0 mm ventral to bregma (DV))^61^. In brief, adult mice >8 weeks old were first anaesthetized with 1.5-3% isoflurane at an oxygen flow rate of 1 L/min. The head fur was shaved, and the incision site was sterilized with 70% ethanol prior to the beginning of the surgical procedure. The mice were then mounted on a stereotaxic device (51730D, Stoelting, IL), and a heat pad (BWT-100A, BRC, Nagoya) was placed underneath the mouse to maintain body temperature at 37 °C. After the scalp was incised and pulled aside, a 2 mm-diameter craniotomy was created with a surgical drill on the skull above the injection site. Through a glass-capillary injection pipette (25 μm inner-diameter tip), 1 μl of AAV5-Syn-GCaMP6f-WPRE-SV40 (titre 2.8 × 10^13^ GC/ml; gift from Douglas Kim & GENIE Project; Addgene viral preparation #100837-AAV5)^62^ was injected using a microinjection pump (Nanoliter 2010, WPI, FL). After the injection, the scalp was sutured and treated with povidone-iodine. DG consists of granule cells (~90%) and contains other types of excitatory neurons, such as mossy cells in hilus and interneurons^47^. GCaMP6f expression is driven by the Syn-promoter; however, it is not cell type specific. Therefore, its expression is expected to be induced not only in granule cells, but also in mossy cells.

#### GRIN lens implantation

At least two weeks after the viral injection, a gradient refractive index (GRIN) lens (Inscopix 1050-002202; 1 mm diameter; 4.0 mm length) was implanted into each mouse. The mouse was mounted on the stereotaxic device as described above, and the scalp was removed. After exposure of the skull and removal of the overlying connective tissue, we made a cranial hole that was slightly larger than the diameter of the GRIN lens. To make a “pre-track” for the GRIN lens insertion, we made an approximately 2 mm-wide incision in the exposed cortex down to 1 mm from the brain surface and then slowly embedded the lens into the dorsal DG (AP: −2.0 mm, ML: +1.0 mm, DV: −1.75 mm). After temporarilly immobilizing the lens with a UV-curable resin (Primefil, Tokuyama Dental, Tokyo), the lens and a stainless frame (CF-10, Narishige, Tokyo) were fixed to the exposed skull using dental cement mixed with black dye (Sudan Black, Sigma, MO). The exposed tip of the GRIN lens was covered with dental silicone (Dent Silicone-V, Shofu, Kyoto). After lens implantation, analgesic and anti-inflammatory agents (Flunixin, 2 mg/kg; Fujita, Tokyo) and antibiotics (Tribrissen, 0.12 ml/kg; Kyoritsu Seiyaku, Tokyo) were injected intraperitoneally. After each experiment, we confirmed that the GRIN lens was implanted in the upper side of the granule cell layer, where most neurons are GCs, by histological analysis (Supplementary Fig. 11).

#### Baseplate attachment

At least two weeks after GRIN lens implantation, the baseplate of a miniature micro-endoscope (nVista 2.0, Inscopix, CA) was attached over the GRIN lens by a conventional method^31^. Briefly, the mice were anaesthetized and mounted onto the stereotaxic device as described above, and a baseplate attached to the miniature microscope was placed on the GRIN lens using Gripper (Inscopix, CA). By monitoring the fluorescent images of GCaMP-expressing neurons, the optimal location was determined (where the largest number of neurons were in focus), and the baseplate was fixed with dental resin cement (Super-Bond C&B, Sun Medical, Shiga) at this position. In cases where we failed to identify neurons at this stage (usually due to the failure of GRIN lens implantation), the baseplate was not attached, and such mice were excluded from the experiment. After baseplate attachment, a baseplate cover was placed on the baseplate until Ca^2+^ imaging was performed.

### Ca^2+^ imaging in freely moving mice

Ca^2+^ imaging of DG neurons was performed while the mice were freely travelling in an open field (OF) or T-maze, each on a different day. Each different group of mice was used for OF or T-maze. Prior to Ca^2+^ imaging, the mice were lightly anaesthetized, and a miniature microscope (nVista 2.0, Inscopix, CA) was mounted onto the baseplate of the mice. The mice were then placed back in the home cages, which were then transferred to a sound-proof behavioural experiment room. At least 30 min after recovery from anaesthesia, the mice were subjected to OF or T-maze tests while the Ca^2+^ signals of their DG neurons were obtained at a 3 Hz sampling rate with 1440 × 1080 pixel resolution. We applied 475/10 nm LED light for the excitation of GCaMP6f fluorescence (approximately 0.24 mW/mm^2^ at the bottom of the GRIN lens).

### Behavioural experiments

Before every behavioural experiment, the OF or T-maze apparatus was cleaned using weakly acidified hypochlorous water (super hypochlorous water; Shimizu Laboratory Supplies, Kyoto, Japan) to prevent bias due to olfactory cues. All behavioural experiments were carried out in a sound-proof room, and the behaviour of the mice was monitored through a computer screen located outside the room to minimize artefactual cues due to the presence of the experimenter. Mouse behaviour was recorded at a 3 Hz sampling rate.

#### Open field test

More than a week after baseplate attachment, mice were habituated to the test environment. Each mouse was lightly anaesthetized, and a dummy camera (Inscopix, CA) was mounted on the mouse. At least 30 min after recovery from anesthesia, mice were placed for 2 hr in the OF arena (40 cm × 40 cm × 30 cm; width, depth, and height, respectively; O’Hara, Japan), made of opaque white plastic. The OF apparatus was evenly illuminated with 100 lux white LED light installed above the apparatus. This habituation session was repeated for three days. One day after the final habituation session, OF experiment was performed. Prior to experiment, the mice were weakly anaesthetized with isoflurane, and an nVista miniature microscope was mounted onto the head stage. The mice were then habituated in the testing room for at least 30 min after recovery from the anaesthesia. Following the habituation in the testing room, each mouse was placed in the OF arena, and neuronal activity was recorded for 30 min. In order to obtain the location time sequence of each mouse, the images of the mouse were automatically processed by an ImageJ plugin (Image OF, freely available on the Mouse Phenotype Database website: http://www.mouse-phenotype.org/software.html).

#### T-maze test

The T-maze test was conducted using an automatic T-maze apparatus (O’Hara, Japan) as previously described^46^. In brief, the maze consists of the stem of the T (13 cm × 24 cm); the left and right (L/R) arms (11.5 cm × 20.5 cm each side); and connecting passageways from the end of the L/R arms to the starting compartment. These compartments are partitioned by sliding doors that open downward. The mice were subjected to a spontaneous alternation protocol for five sessions, with at least 1 day (2 days maximum) of rest between sessions. Each session consisted of 10 trials with a 50-min cut-off time, and each trial consisted of a first and second run. On the first run, the mouse was forced to choose one of the L/R arms (forced choice). After the mouse was in the L/R arm for more than 10 sec, the door to the connecting passageway was opened, which allowed the mouse to return to the starting compartment. When the mouse returned to the starting compartment, all the doors of this component were closed; then, after 3 sec, the doors connecting to the L/R arms opened for the second run, in which the mouse could freely choose either of the L/R arms (free choice). The percentage of trials in which the mouse entered the arm opposite to their forced-choice run (the “correct” arm) was calculated. The choice of the L/R arm for the forced trials was varied pseudo-randomly across trials using a Gellermann schedule so that the mice received equal numbers of left and right presentations. Data acquisition, control of the sliding doors, and data analysis were performed with ImageTM software (freely available on the Mouse Phenotype Database website: http://www.mouse-phenotype.org/software.html)

### Detection of Ca^2+^ transients

To extract the activity patterns of individual DG neurons from the obtained fluorescent images, we used Inscopix Data Processing Software (IDPS). Briefly, the data files of the raw sequential fluorescent images obtained by nVista (Inscopix) were imported to IDPS, and motion correction was applied. The instantaneous fluorescence of the motion-corrected images was then normalized by its average fluorescence over the entire recording period, producing fluorescence change ratio (ΔF/F) images. Then, individual DG neurons were identified by automated PCA/ICA (PCA, principal component analysis; ICA, independent component analysis) segmentation of activity traces. For PCA/ICA segmentation, default parameters, whose values were confirmed to work well across most of the neuronal activity patterns in the cortex and CA1 of the hippocampus, were used (number of ICs, 120; number of PCs, 150; ICA max iterations, 100; ICA random seed, 0; ICA convergence threshold, 0.00001; block size, 1000; ICA unmixing dimension, spatial; ICA temporal weights, 0.00; arbitrary units within IDPS). After cell identification, longitudinal cell registration was performed to align cell maps across multiple sessions as necessary. Finally, the temporal pattern of each the timings of DG neuron’s activity was identified by an event detection algorithm available in IDPS (event threshold factor, 4 median absolute deviations; event smallest decay time, 0.2 sec) and the temporal pattern of activity was expressed as a time series of binarized signals for further analysis.

### Estimation of the active population of DG neurons

Estimation of the active population of DG neurons was performed on the basis of the instructions of Dr. Jonathan Zapata, Inscopix Inc.. In brief, after motion correction of the raw fluorescent images from the maximum intensity projection of GCaMP fluorescence images, the total number of neurons in a field of view was manually counted. Likewise, the maximum intensity projection of the fluorescence change ratio (ΔF/F) images was generated, and the total number of active neurons during the 30 min recording was manually counted. The maximum projections of GCaMP fluorescence images and ΔF/F images were calculated throughout all frames in the 30 min recording in the OF using the “movie projection” tool available in IDPS and saved as tiff files. We counted the number of cells in the maximum projection image frames using the multipoint tool of ImageJ (available from https://imagej.nih.gov/ij/). To confirm the activity of all the detected cells, we manually drew regions of interest (ROIs) over all the cells in the ΔF/F projection images and confirmed the traces of the Ca^2+^ signals with IDPS. All neurons that exhibited calcium transients individually in the soma at least once during the entire recording period were considered “active”. All the image data that were used to perform active cell counting and the detailed method are available at “SSBD:repository” database [http://ssbd.qbic.riken.jp/set/20200603/].

### Definitions of the position, speed, and motion direction of mice

i. Position: The OF arena (which had an area of 40 cm × 40 cm) was represented as 200 × 200 pixel grid. The position of the mouse is determined from the centroid of its shadow on the camera. We then assigned a label corresponding to the discrete location of the mouse (e.g., [10, 100]) to each time bin (=1/3 sec).
ii. Speed: From the distance travelled between 1 sec before and after a given time point, we calculated the speed of mouse at that moment, and assigned this speed (cm/sec) to each time bin.
iii. Motion direction: The visual tracking system that we used does not allow direct measurement of head direction. Instead, we indirectly estimated the direction of motion from the changes in the position of the mouse. The motion direction (in radians) was computed from the direction of change in two subsequent mouse positions (1 sec before and after a given time point) in the x-y plane and assigned this to each time bin. North was defined as 0 radians; west was defined from 0 to π radians, and east was defined from 0 to −π radians.

### Statistical analysis of spatial, speed and direction information

To quantify the tuning specificities of neurons with position, speed, and motion direction, we measured their specificity in terms of the information rate of cell activity and defined them as (i) spatial, (ii) speed, and (iii) direction information^44^. The Ca^2+^ event rate in Ca^2+^ imaging is considerably lower than in electrophysiological recordings (on average, approximately 30 Ca^2+^ transients per 30 min session). If the discretization of position, speed, and motion direction it too fine for the number of events in the recorded cells, we will not be able to obtain a proper null distribution when creating the shuffle data for that cell. Therefore, we set the resolution of the discretization of position and speed to be lower than those commonly performed.

i. We used a 2×2 square grid for measurement of spatial information and computed the amount of Shannon information that a single Ca^2+^ transient conveyed about the animal’s position. The spatial information *I* (bits per Ca^2+^ transient) of a cell was calculated as the mutual information score between the occurrence of a single Ca^2+^ transient of the cell and the animal’s behavioural state of position using the formula:

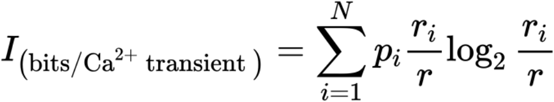

where *i* is the bin number corresponding to the physical parameter (in this case, spatial position in the OF; i = 1–4 from the 2×2 square grid), *N* is the total number of bins, *p* is the probability that the mouse occupied bin *i*, *r_i_* is the mean transient rate at bin *i*, and *r* is the overall mean transient rate.
ii. To measure speed information, we applied the same formula to the speed after discretizing it to a binary state: Run (for speeds >1 cm/sec) and Stop (for speeds <1 cm/sec).
iii. Similarly, we measured direction information by applying the same formula to the motion direction after discretizing the full angle to 8 bins of 45 degrees each.

### LR indices of neurons in the T-maze test

To quantify the left- or right-preference of each neuron’s activity for the current and future location in the T-maze, we generated a parameter called the LR index. For each neuron, we measured the average Ca^2+^ transient rates during the forced arm and decision periods for the left- and right-choice trials—that is, when the mouse is in the left or right arm of the T-maze, respectively—and computed the LR index as follows:

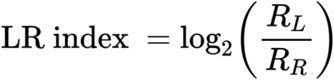

where *R_L_* is the average Ca^2+^ transient rate across all left-choice trials, and *R_R_* is that from the right-choice trials. All neurons are assigned an LR index in both the forced arm and decision periods. A positive value indicates a neuron’s preference for the left arm, whereas a negative value indicates a neuron’s preference for the right arm.

### Data Shuffling

To assess the statistical significance of information coding of individual neurons, we computed chance distributions of the shuffled data using two common methods previously described in the literature^32,18,9^.

Random permutation permutates calcium events (Supplementary Figure 12). We divided the calcium event data into 1000 segments along the time axis and randomly sorted them to generate permutated data of calcium events. This method destroys temporal structures of neural activity and temporal correlations between neural activity and behavioral variables (e.g., position, speed, and motion direction in open field test); however, the overall neural activity is maintained across cells. We repeated this procedure 1000 times to obtain the distribution of 1000 shuffled data. For single-cell statistics, we compared the original information of individual cells with the null distributions of 1000 values of shuffled data generated from the original cell (Supplementary Figure 2). If the original value of information of a cell exceeded 3 sigma from the shuffled distribution, the cells were defined as carrying significant amounts of information. For group comparison (for example, Obs. vs Shuff. in Fig. 1d, 4c, and 4d), we pooled all the shuffled data in a group together, and distributions of the original cells were compared with null distributions of the shuffled data.

Random scrambling is a method that maintains temporal dynamics of neural activity data while disrupting the relationship with behavioral patterns, (for example, the calcium event timeseries and the animal’s position (Supplementary Figure 12)). We shift the whole vector of the calcium event time series in time by a random amount in a torus, that is, points that went beyond the time limits of the data were reinsert from the other side. This procedure disrupts the relationship between neural activity and animal behaviour, but preserves the temporal patterns of these variables. By repeating this procedure while changing the number of frames to be shifted at random, we obtain the null distributions of shuffled data (Supplementary Figure 12). Information statistics are performed in the same way as the first shuffling method.

Since we found that the information statistics and decoding results of the shuffled data did not differ significantly between these shuffling strategies (Supplementary Figure 12), we adopted the random permutation method for the generation of shuffled data.

### Decoding position, speed, and motion direction in the open field

To determine the manner in which the OF behavioural parameters are encoded in the DG, we trained decoders with machine learning methods to separately predict position, speed, and motion direction from the population Ca^2+^ activity. We assigned the labels of the discretized behavioural parameters of the mouse (position (cm), speed (cm/sec), and motion direction (−π – +π radians)) and the binary values of the Ca^2+^ signal (0 or 1) of all neurons to each time bin. We then divided the Ca^2+^ imaging data and behavioural data from each 30 min trial into the first 15 min and last 15 min halves, which were designated training and test data, respectively. For each pair of behavioural parameter (position, speed, or motion direction) and the value of the Ca^2+^ signal in the training data, we trained the decoders with one of eight different machine learning methods (Dense Feedforward Neural Network (DNN), Gated Recurrent Unit (GRU), Long Short-Term Memory (LSTM) network, Recurrent Neural Network (RNN), Support Vector Regression (SVR), Wiener Cascade (WC), Wiener Filter (WF), Extreme Gradient Boosting (XGB); the codes were obtained and modified from Glaser *et al*., arXiv, 2018^63^). The decoding accuracy of the position/motion direction is reported as the mean absolute error in the distance between the predicted and actual position/motion direction. In the case of speed, decoding accuracy was reported as the correlation between the predicted and actual instantaneous speed. This is because speed, unlike position and motion direction, is not limited to a certain range and mean absolute error of speed may depends on the average locomotion speed of each individual mouse, which would be inappropriate for evaluation of decoding error. To assess the statistical significance of the decoding accuracies, the decoding error from the observed data was compared with that of the shuffled data, which is created by dividing the Ca^2+^ imaging data into 1,000 segments and sorting them randomly as described above. Furthermore, we compared the decoding accuracies of the three behavioural parameters among the 8 decoders and found that they were not significantly different (Supplementary Fig. 3). Consequently, we reported the decoding results that were obtained with the LSTM Network, which showed slightly better decoding performance in wild-type mice than others. For decoding position and motion direction, we identified and removed time bins when the mouse moved at speeds below 1.0 cm/sec and 4.0 cm/sec, respectively, to obtain optimal decoding results. Details on the methods used in thresholding the data according to the movement speeds of the mice are described in the Supplementary Notes and Supplementary Fig. 10.

### Decoding current and future location in the T-maze test

We similarly sought to decode the left or right preferences for the current and future locations in the T-Maze test using the population Ca^2+^ activity with machine learning methods. For each trial of the forced arm and decision periods, we assigned a label corresponding to the left or right choice of the mouse and the average Ca^2+^ transient rate of each neuron during the period. We then used a support vector machine (SVM) classifier function in MATLAB for binary classification of the left or right decision (MathWorks, MA). All 50 trials from each mouse were used for 25-fold cross-validation; the 50 trials were randomly divided into 48 trials of training data and 2 trials of test data. We repeatedly trained the binary classifier using randomly selected training data and performed left or right predictions for the remaining test data to evaluate the decoding accuracy. We also shuffled the Ca^2+^ transient data and performed the same decoding analysis. As with the open field behavioural parameters, the decoding accuracy was compared between the shuffled and unshuffled data.

### Population vector overlap (PVO)

To quantify the similarities in population activity patterns of neurons between different locations in the DG, we calculated the population vector overlap (PVO)^64^. The PVO for a population of N neurons in two different conditions (x, y) was defined as

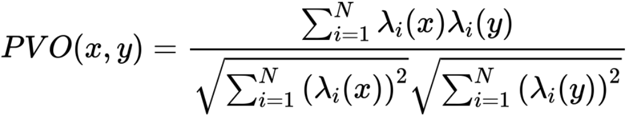

where *i* is the neuron number; λ_j_(x) and λ_j_(y) are the average Ca^2+^ transient rates of neuron j in conditions x and y, respectively; and *N* is the total number of neurons. For a given population of neurons and a pair of conditions, a lower PVO indicates that the activity patterns of the neurons are distinct between the two conditions.

### Code and data availability

The data analysis was performed using custom code written in MATLAB (MathWorks, R2018a) and Python (Version 2.7.12). For decoding analysis by machine learning, we obtained Python code available at https://github.com/KordingLab/Neural_Decoding (used in Glaser *et al*., arXiv, 2018^63^) and modified them for our purpose. Raw imaging data in this study is uploaded at Systems Science Biological Dynamics repository (SSBD:repository; http://ssbd.qbic.riken.jp/set/20200603). Processed imaging data and behavioural data in this study are available at https://github.com/tmurano. Codes will be made available upon reasonable request.

## Supporting information

Demonstration_OF_decoding_wikd-type

Demonstration_OF_decoding_aCaMKII

## Supplementary Notes

### Velocity filtering for position and direction decoding (Supplementary Fig. 10)

The activation of hippocampal place cells during immobile periods, which occurs in conjunction with hippocampal sharp wave ripples (SWRs), is known not to represent the current position of animals^65,66^. Therefore, in analyses of position information in the hippocampus, periods of immobility are generally excluded from the data. We did not know whether immobile periods should also be excluded for data obtained from the dDG, as it is not known whether neural activity equivalent to SWRs can be observed in the Ca^2+^ imaging in this region. Therefore, to test the validity of this procedure, we examined whether removing periods of immobility would improve the accuracy of position decoding in the dDG (Supplementary Fig. 10, left panel). We removed the time bins in which the movement speed of the mouse was below the threshold (from 0.0 cm/sec to 3.0 cm/sec) and performed position decoding using the remaining data. In wild-type mice, removing time bins in which the speed was below 1.0–1.25 cm/sec reduced the position decoding error (Supplementary Fig. 10, left panel). These results suggest that neural activity during periods of immobility that do not represent current location may also be observed in the Ca^2+^ imaging data of the dDG. Based on these results, all periods in which the mouse’s speed was below 1 cm/sec were eliminated from the analysis of position information. (This threshold is roughly equivalent to those commonly used in hippocampal CA cells (1.0-2.0 cm/sec)^32^).

The motion direction of the mouse was estimated from the changes in the mouse position in the open field. Even small movements such as grooming, rearing or head turning were calculated as actual mouse movements. To exclude these periods and more accurately measure motion direction, we examined the speed threshold below which movements could to be removed from the datasets (Supplementary Fig. 10, right panel). We removed all the time bins in which the speed of the mouse was below the threshold (from 0.0 cm/sec to 10.0 cm/sec) and performed direction decoding using the remaining data. In wild-type mice, the decoding error for motion direction decreased after applying a threshold of 4.0-8.0 cm/sec to remove movement periods, suggesting that small movement periods below these thresholds were useless for accurate direction decoding (Supplementary Fig. 10, right panel). Based on these results, all periods when the mouse ran slower than 4.0 cm/sec were removed from our datasets for the analysis of motion direction.

## Acknowledgements

We thank Wakako Hasegawa, Yumiko Mobayashi, Misako Murai, Tamaki Murakami, Miwa Takeuchi, Yoko Kagami, Harumi Mitsuya, Yoshihiro Takamiya, and other members of the Miyakawa laboratory for their support. We also thank Jonathan Zapata (Inscopix) for his detailed advices on the image analysis. This work was supported by a JSPS Grant-in-Aid for Scientific Research on Innovative Areas (grant #JP16H06462), the AMED Strategic Research Program for Brain Science (grant #JP18dm0107101), and a grant from Astellas Pharma Inc.

## Author Contribution Statement

T. Murano A. Nakao, R. Nakajima, J. Yamamoto, and T. Miyakawa designed the experiments and analyses. A. Nakao, N. Hirata, and R. Nakajima performed the behavioural testing and Ca^2+^ imaging. T. Murano, R. Nakajima, S. Amemori, A. Murakami, and Y. Kamitani analysed the data. T. Murano, R. Nakajima, J. Yamamoto, and T. Miyakawa prepared figures and wrote the manuscript. T. Miyakawa supervised all aspects of the present study. All authors reviewed the manuscript.

## Additional Information

Processed imaging and behavioural data are available at https://github.com/tmurano. Raw images and behavioural data are available at http://ssbd.qbic.riken.jp/set/20200603/.

Correspondence and requests for materials should be addressed to T.M.

