## Supplementary Figures for "Multiple types of navigational information are independently encoded in the population activities of the dentate gyrus neurons"

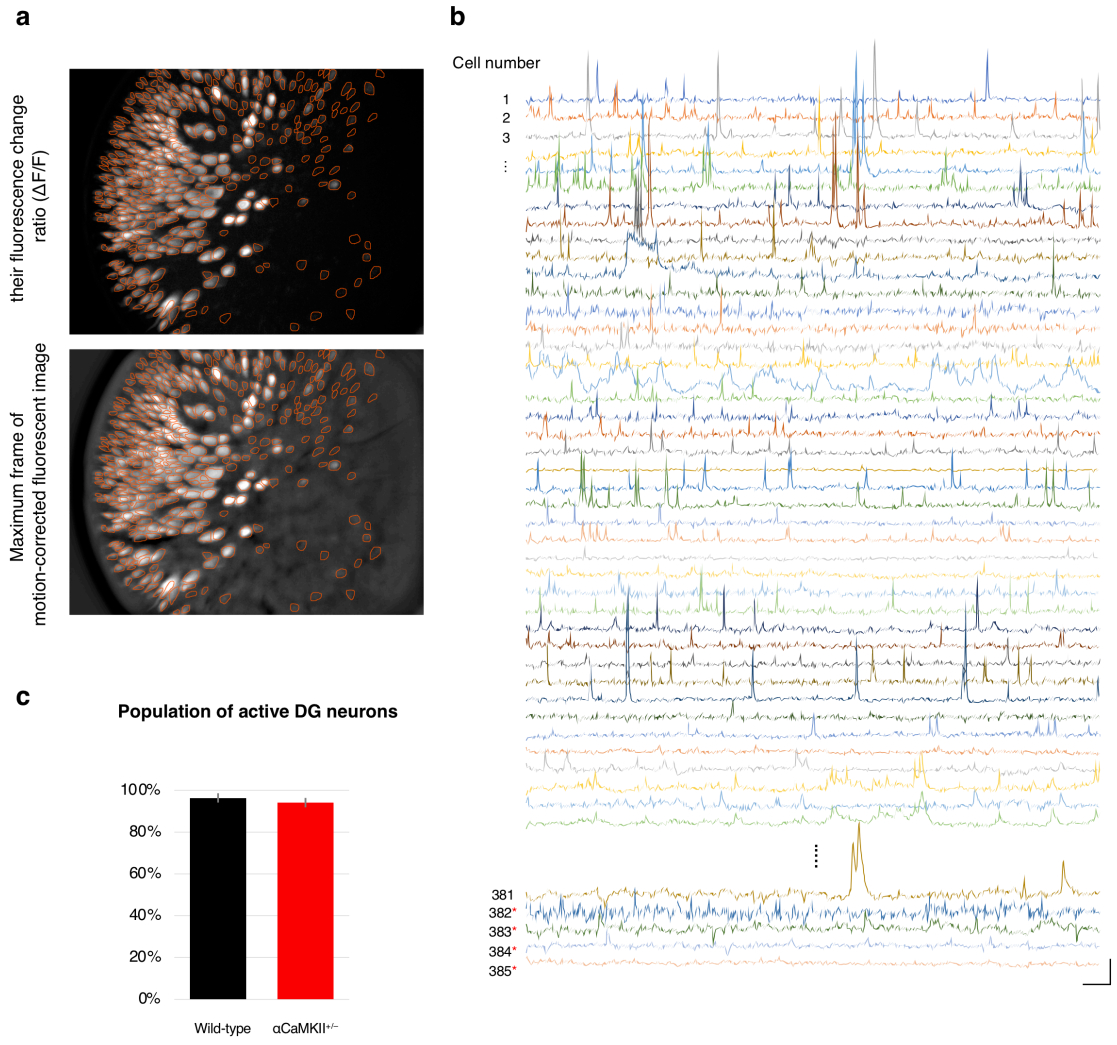


***Supplementary Figure 1. Proportion of active DG neurons.***

**a,** Fluorescence change ratio (ΔF/F) image (upper panel) and maximum frame of motion-corrected (MC) fluorescent image (lower panel) of a representative wild-type mouse. Orange circles indicate the ROI of cells from the maximum projection image.

**b,** Representative $Ca^{2+}$ signals (ΔF/F) of individual dDG neurons, indicated by the regions of interest (ROIs) in (a). Fifty representative cells of a total of 384 cells are shown, and red asterisks indicate GCaMP-expressing neurons whose activity was not detected in the 30 min recording. Scale bars: 30 sec (horizontal) and 5% ΔF/F (vertical).

**c,** The proportion of dDG neurons in the field of view that were active. The number of neurons demonstrating GCaMP expression was manually counted from the frame with maximum MC fluorescence, and the number of neurons with calcium transients (active neurons) was counted from their fluorescence change ratio ($\Delta F/F$) images. Bar graphs show the average proportion of active dDG neurons. Error bars indicate SEM.

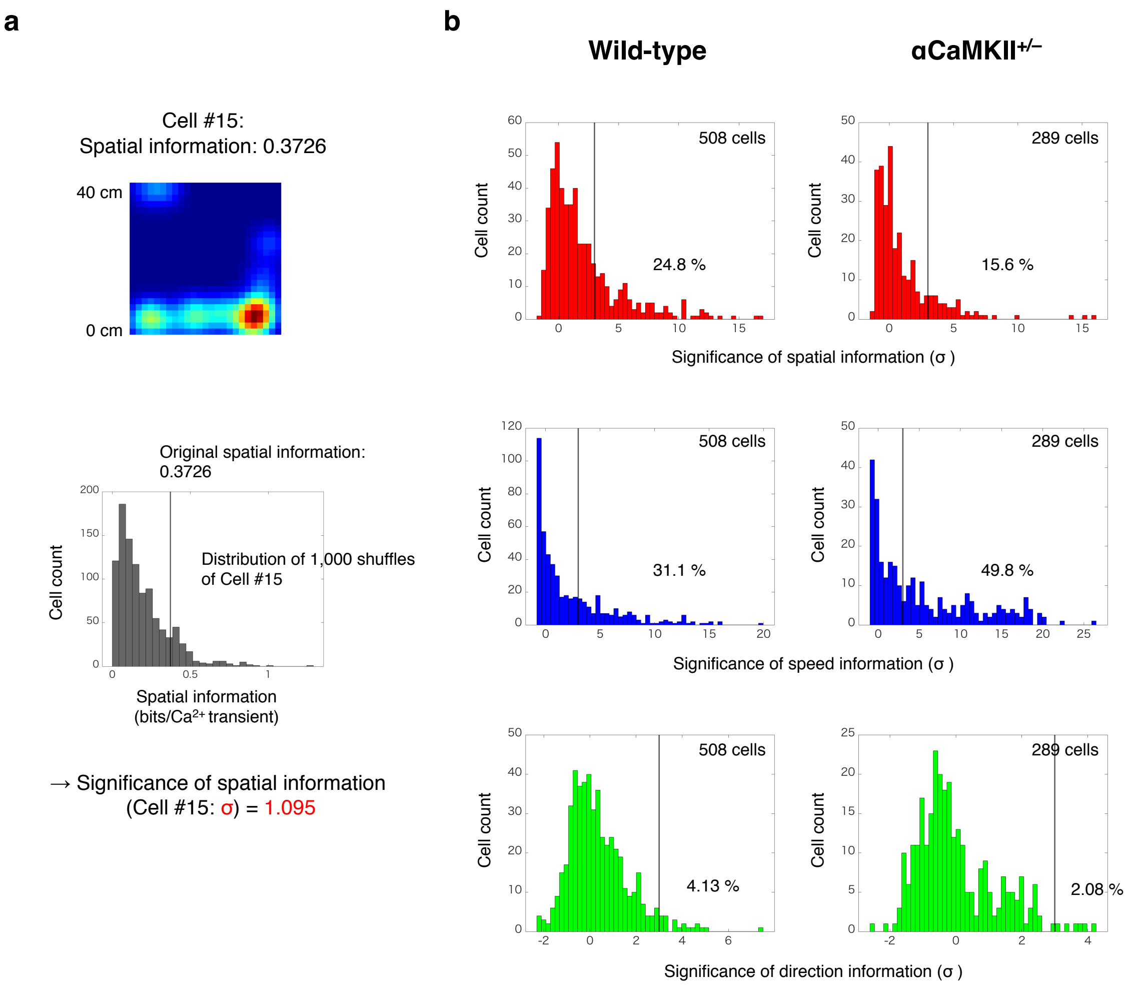


***Supplementary Figure 2. Significance of spatial, speed, and direction information of each dDG neuron.***

**a,** Top: Colour-coded spatial tuning map of representative dDG neuron (Cell #15). Bottom: Distribution of spatial information from 1,000 shuffled data generated from Cell #15. The vertical black line shows the original spatial information for Cell #15. Significance of spatial information of the cell is calculated from the value of the original spatial information in the normalized distribution of shuffled data.

**b,** Distribution of significance of spatial, speed, and direction information for all neurons from all animals (508 neurons from 7 wild-type and 289 neurons from 5 αCaMKII$^{+/-}$ mice). The vertical black line indicates where the significance of information content (σ) is 3, and the numbers on the lines represent the percentage of neurons exceeding this value.

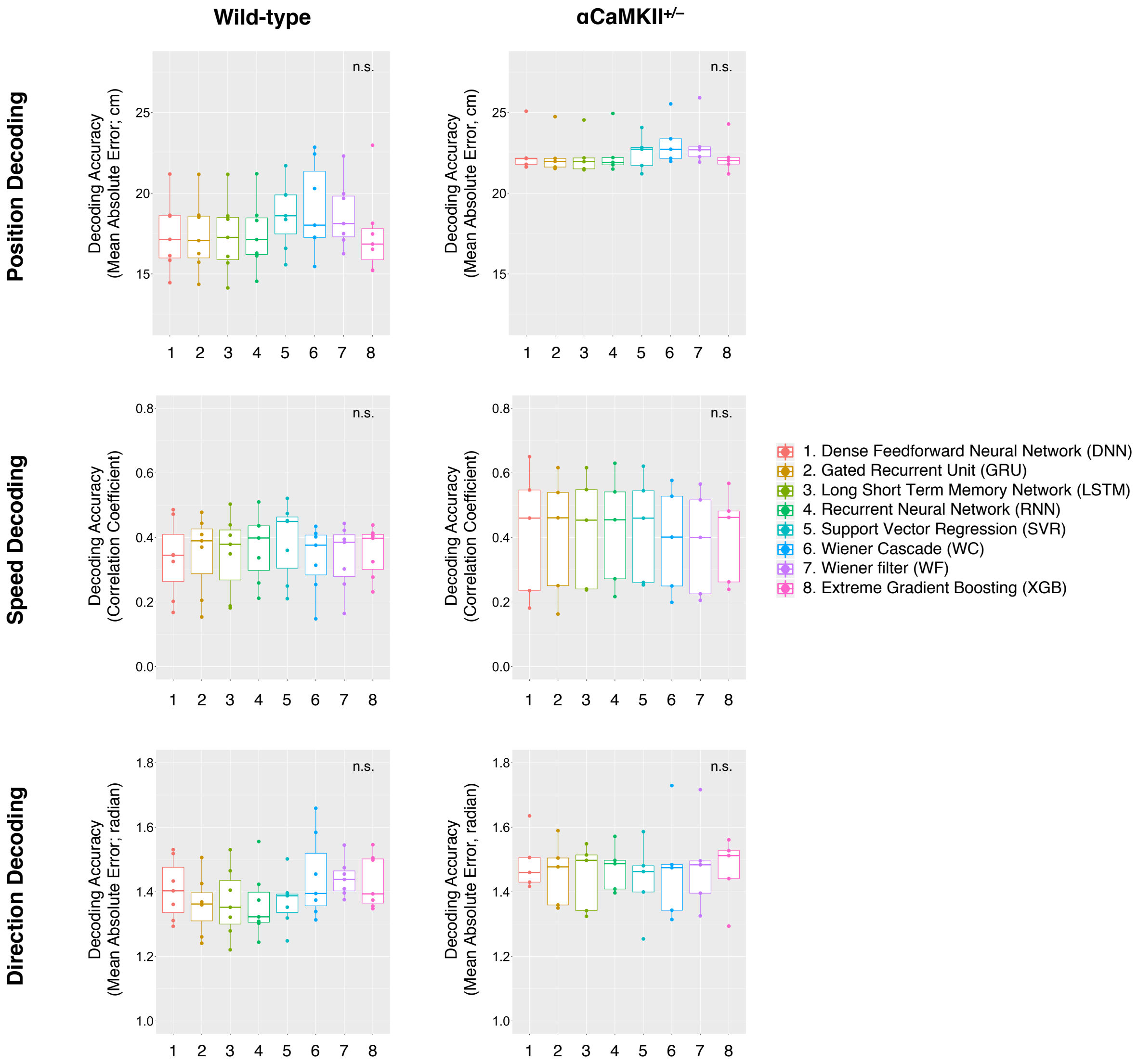


***Supplementary Figure 3. Machine learning methods used to decode position, speed, and motion direction.***

We tested 8 different machine learning methods for separately decoding the position, speed, and motion direction in mice in the open field test (as listed in the figure). Overall, there was no significant difference in the decoding accuracies among these models.

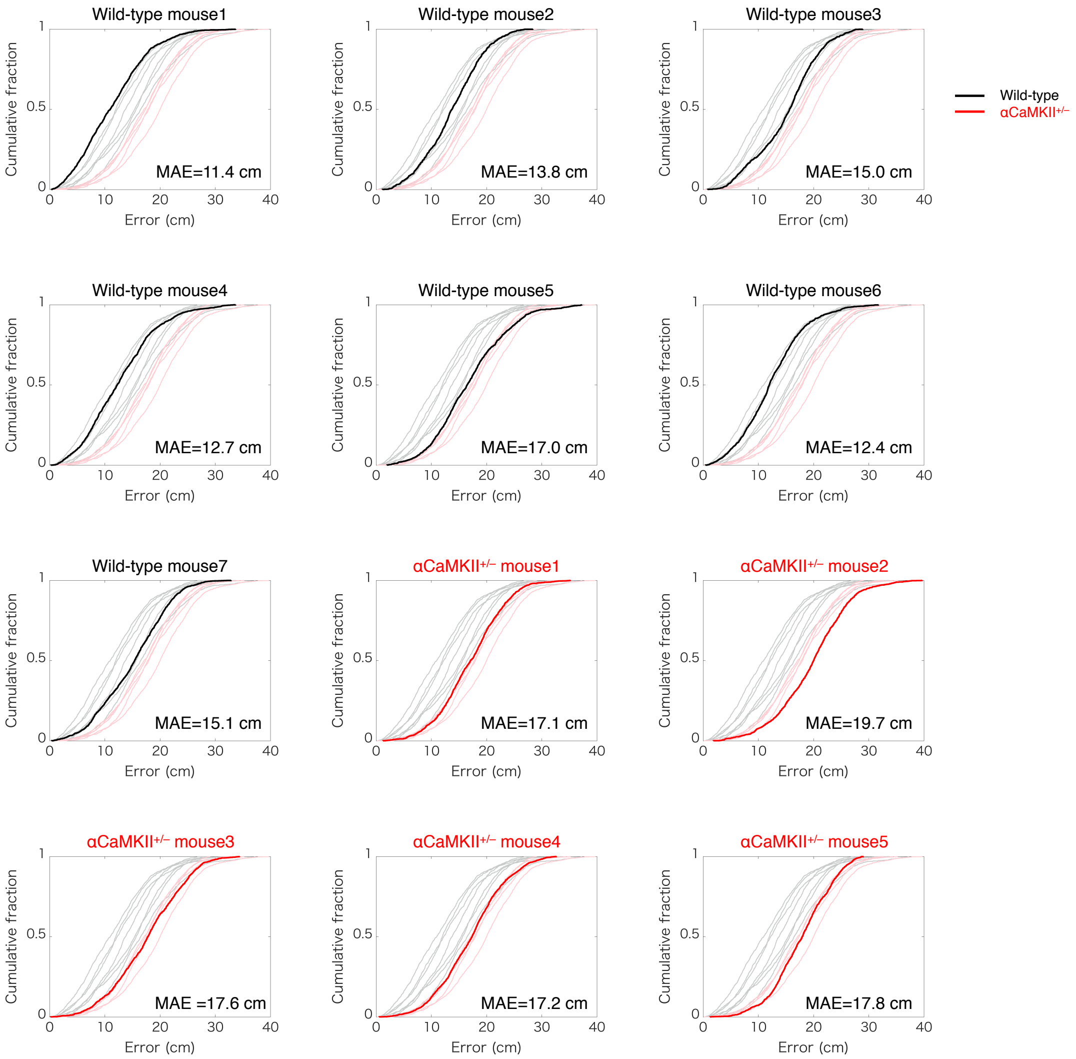


***Supplementary Figure 4. Position decoding error in the open field***

Cumulative distribution functions (CDFs) of the position decoding error. The results from all the mice (black; 7 wild-type mice, red; 5 αCaMKII$^{+/-}$ mice) are shown. Mean absolute errors (MAE, same as in Figure 2d) of each mouse are also shown in each figure. Note that all of the decoding errors of wild-type mice were smaller than those of αCaMKII$^{+/-}$mice.

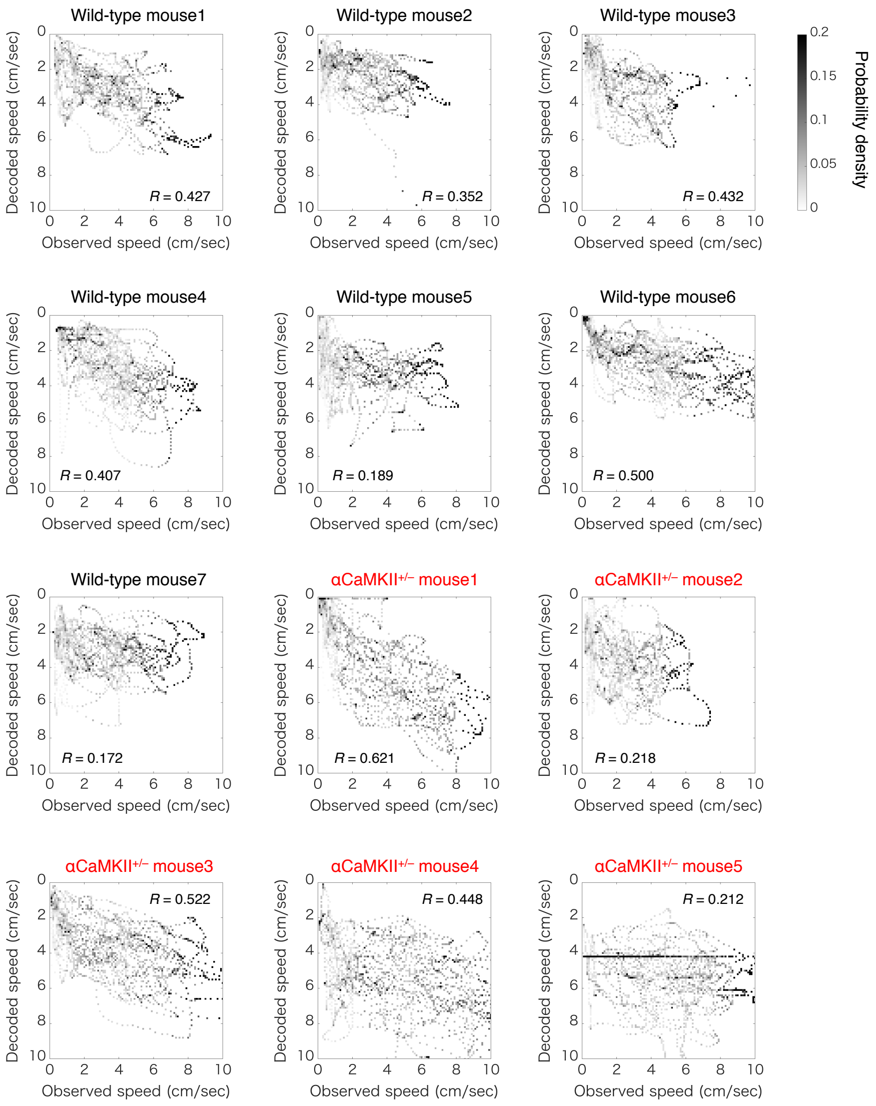


***Supplementary Figure 5. Speed decoding accuracy in the open field***

Confusion matrices for speed decoding results for all wild-type and αCaMKII$^{+/-}$ mice. The x-axis represents the actual mouse speed, and the y-axis represents the estimated mouse speed. Each column corresponds to the estimated probability distribution over speed for a given true speed (0.1 cm/sec bins). The correlation coefficient (*R*) between the observed and decoded speeds (same as in Figure 2e) is shown in each mouse.

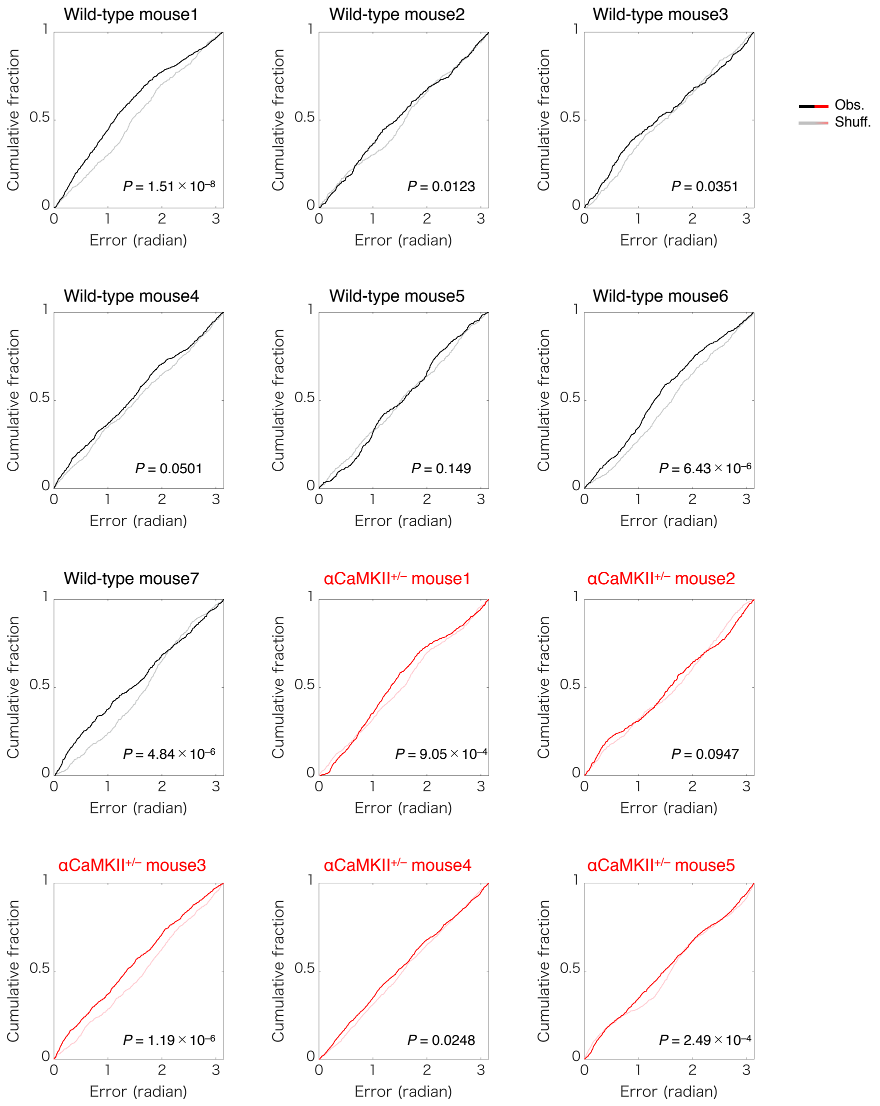


***Supplementary Figure 6. Direction decoding error in the open field***

Cumulative distribution functions (CDFs) of the direction decoding error. The results from all the mice (black; 7 wild-type mice, red; 5 αCaMKII$^{+/-}$ mice) are shown. Mean absolute errors (MAE, same as in Figure 2f) of each mouse are also shown in each figure. The *P*-value in each panel indicates the difference between the distribution of decoding error from observed and shuffled data (Kolmogorov-Smirnov test).

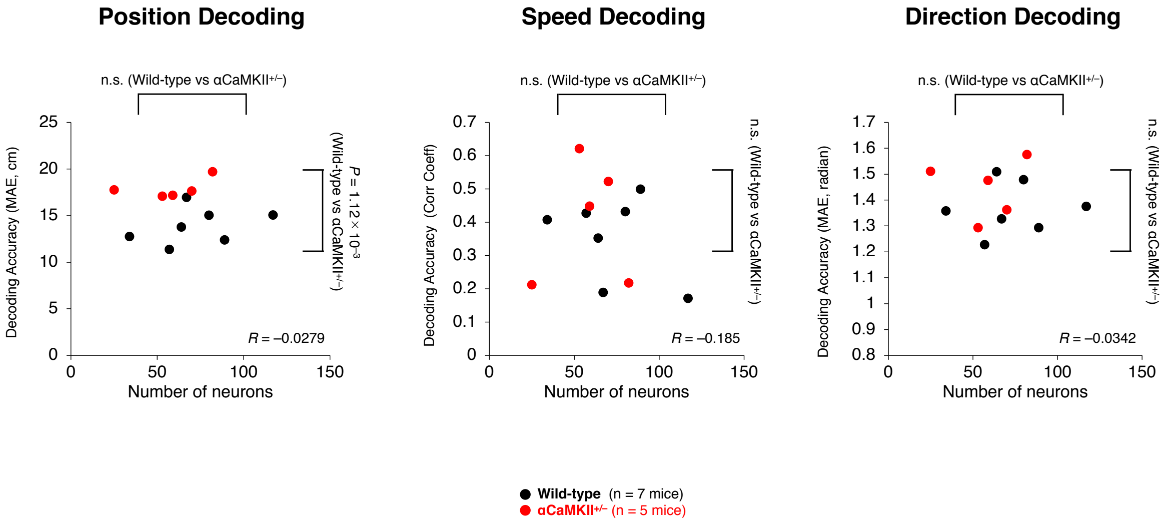


***Supplementary Figure 7. Decoding accuracies for position, speed, and motion direction are less sensitive to the numbers of neurons used for decoding.***

Scatter plots showing the correlation between the decoding accuracies for position, speed, and motion direction and the number of the neurons used for decoding. Each black and red dot corresponds to individual wild-type and αCaMKII$^{+/-}$ mice, respectively. *R* values are correlation coefficients.

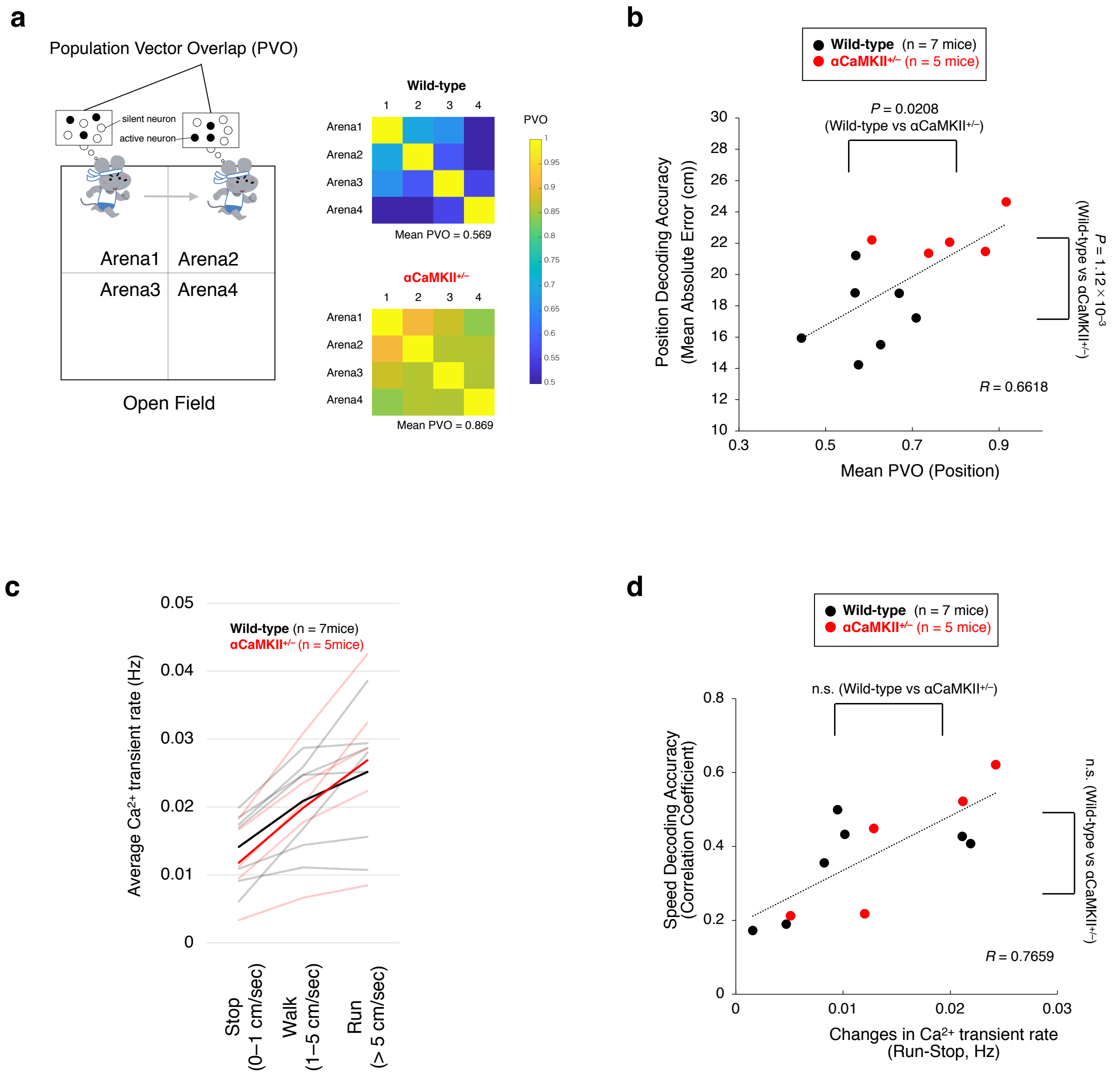


***Supplementary Figure 8. Position and speed decoding accuracies are correlated with the distinctness of neuronal activity patterns and activity frequency, respectively.***

**a,** Similarities in the population activity patterns of dDG neurons among different areas in the open field. The open field is divided into 4 subareas (Arena 1–4), and the population activity patterns of dDG neurons are compared among these subareas. The similarities among the neural activity patterns are evaluated by the population vector overlap (PVO; for details, see Materials and Methods section) of the columns of the average $Ca^{2+}$ transient rate across dDG neurons between two subareas. Colour-coded PVO matrices and their mean PVOs are shown for a representative mouse from the wild-type and $\alpha CaMKII^{+/-}$ groups. A larger PVO indicates higher similarities in the neural activity patterns, and a smaller PVO shows that there are distinct neural activity patterns among subareas.

**b,** Scatter plot showing the correlation between the mean value of the PVO among all subareas and the position decoding accuracy for each mouse. Each black and red dot corresponds to an individual wild-type and αCaMKII$^{+/-}$ mouse, respectively.

**c,** Changes in the average $Ca^{2+}$ transient rate with mouse acceleration. The vertical axis shows the average $Ca^{2+}$ transient rates during the Stop (<1 cm/sec), Walk (1–5 cm/sec), and Run (>5 cm/sec) periods. Each grey and pale red line indicates the average $Ca^{2+}$ transient rates of an individual wild-type and αCaMKII$^{+/-}$ mouse, respectively. The black and red lines show the average of all mice in each group.

**d,** Scatter plot showing the correlation between the change in the $Ca^{2+}$ transient rate (from the Stop to Run periods) and speed decoding accuracy. Each black and red dot corresponds to an individual wild-type or αCaMKII$^{+/-}$ mouse, respectively.

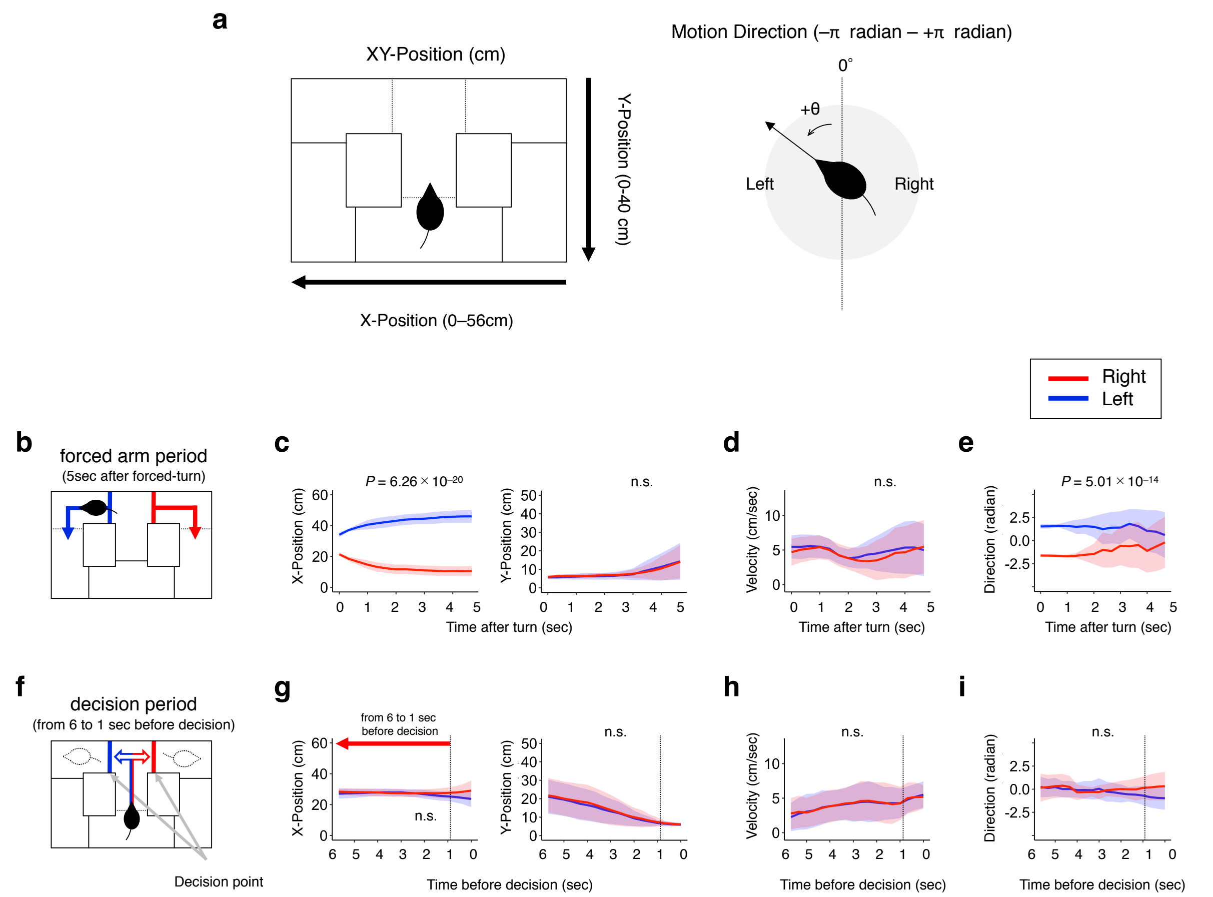


***Supplementary Figure 9. X and Y position, speed, and motion direction of mice during the forced arm and decision periods in the T-maze.***

**a,** Definition of the X position (0–56 cm), Y position (0–40 cm) and motion direction (from –π to +π radians) for mice in the T-maze

**b,** The arm period is defined as the time period 5 sec after the forced turn.

**c-e,** Plots of the mean±SEM X and Y positions (c), speed (d), and motion direction (e) of the mice across all trials for the left (blue lines) and right (red lines) forced choices. The X position and motion direction in the arm period are significantly different between the left and right forced choices.

**f,** The decision period is defined as the time period from 6 to 1 sec before the decision.

**g-i,** Plots of the mean±SEM X and Y positions (g), speed (h), and motion direction (i) of the mice across all trials for the left (blue lines) or right (red) free choice. The X-position, Y position, and motion direction in the decision period are not significantly different between the left and right choices.

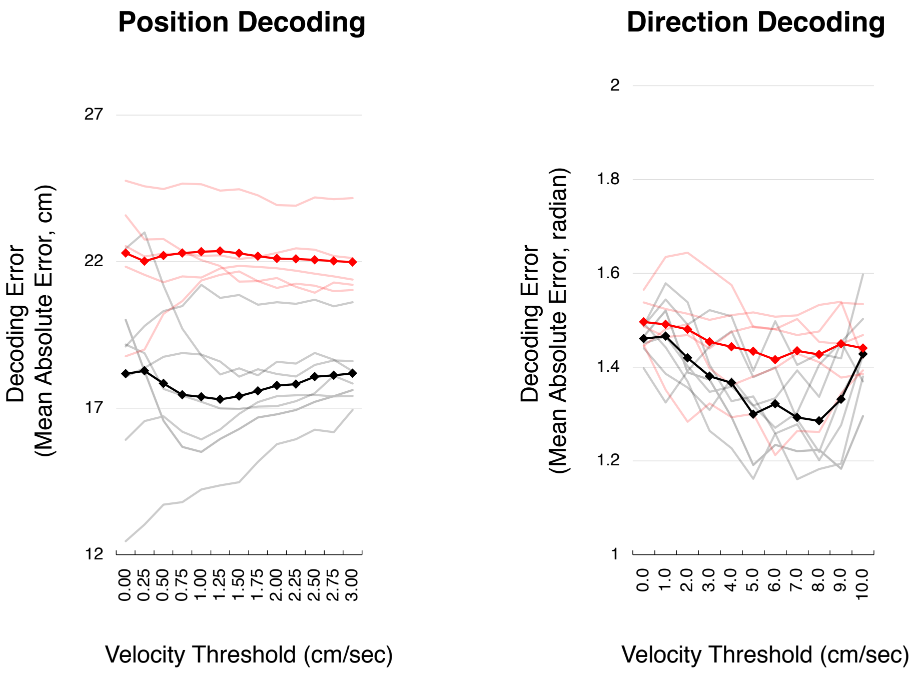


***Supplementary Figure 10. Velocity threshold and decoding accuracy for position and motion direction.***

The horizontal axis indicates the threshold for velocity filtering (for details, see Supplementary Notes). The vertical axis indicates the position decoding error (left panel) and direction decoding error (right panel). The grey and pale red lines indicate individual wild-type and αCaMKII$^{+/-}$ mice, respectively. The black and red lines show the average of all mice in each group.

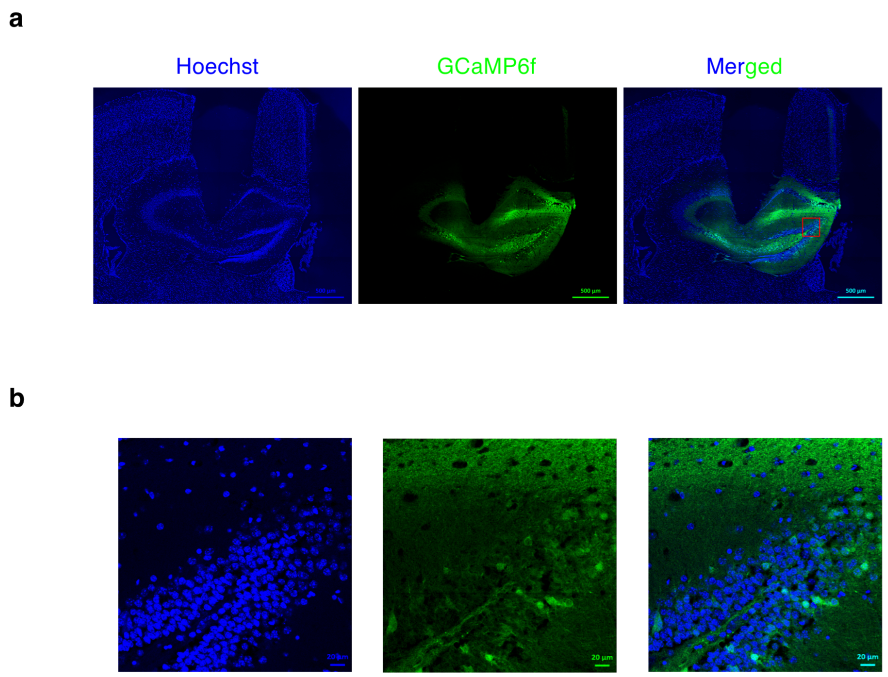


***Supplementary Figure 11. Placement of the GRIN lens implanted in the dentate gyrus.***

**a,** Coronal slice of mouse brain showing the position of implanted GRIN lens. (left: Hoechst; middle: GCaMP6f; right: merged image).

**b,** Enlarged images of the area enclosed by the red rectangle in the panel (a).

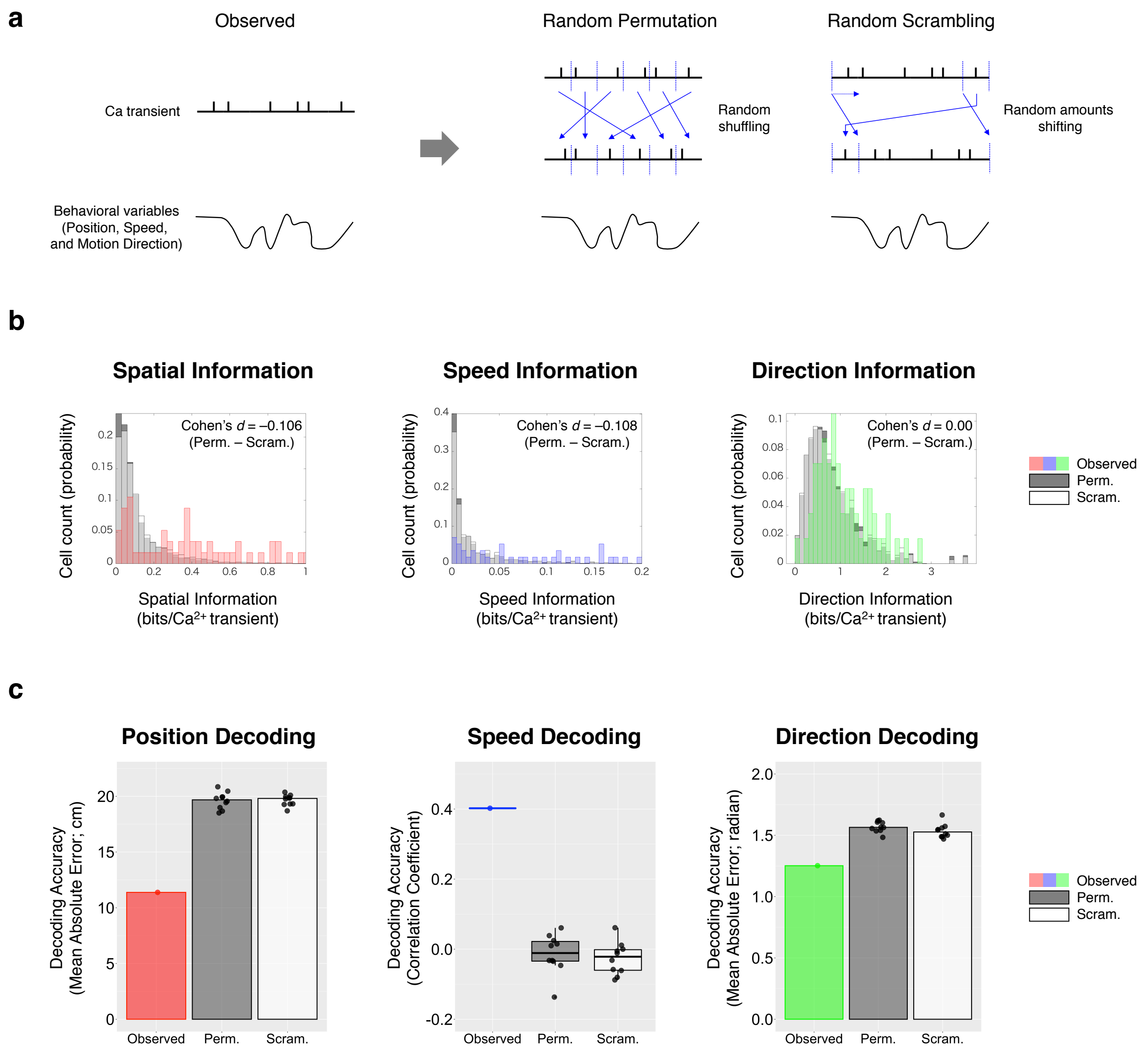


***Supplementary Figure 12.*** Different shuffling methods do not present significant differences regarding information statistics and decoding performance.

**a,** Different methods for generating shuffled data. The random permutation method divided the data into 1000 segments and randomly sorted them, resulting in shuffled data. Thus, both the correspondence between neural activity and behavioral data in the original data, as well as the dynamics of neural activity itself, is impaired. The random scrambling method shifts the time series of the neural activity by a random number of frames while maintaining the temporal dynamics of the neural activity. This method results in disrupted correspondence between the time series of neural activity and behavior; however, the temporal dynamics of behavior and neural activity is preserved.

**b,** Distribution of spatial, speed, and direction information from observed neurons ("Obs.", color-coded bins), as well as for shuffled data ("Perm.", dark gray) and scrambled data ("Scram.", white), for a representative wild-type mouse. We repeated random permutation and random scrambling 1000 times each to generate shuffled data, and there was no significant difference between their distributions.

**c,** Decoding accuracy of position, speed, and motion direction by different shuffling methods. Random permutations and random scrambles were repeated 10 times each to generate shuffled data from a representative wild-type mouse. Decoding analysis was performed on these shuffled data, in which the accuracy was compared, and there was no significant difference between them.

(11 figures)
